# Targeted RNA editing by direct delivery of an adenosine deaminase-antisense oligo conjugate

**DOI:** 10.1101/2025.07.11.664364

**Authors:** Tanner W. Eggert, Dhruv Y. Dhingani, Ralph E. Kleiner

## Abstract

Programmable RNA editing is a promising therapeutic strategy for correcting disease-causing mutations on mRNA, but current approaches rely primarily upon endogenous RNA editing enzymes (i.e. ADARs) that have restricted substrate scope and efficiency. Here, we demonstrate programmable RNA editing with evolved TadA-derived deaminases and 2’-methoxyethyl (MOE)-modified antisense oligonucleotides (ASO) to guide site-directed A-to-I editing. In contrast to ADAR enzymes, TadA proteins modify single-stranded RNA (ssRNA). We profile ASO-guided TadA-based editors on endogenous and disease-relevant mRNAs and develop a “bulge-forming” ASO architecture to constrain RNA editing to the target site. Further, we demonstrate that a covalent adenosine deaminase-ASO “RNP” conjugate formed in the test tube and delivered by lipofection achieves targeted and efficient RNA editing with dramatically lower transcriptome-wide off-target editing as compared to ectopically expressed RNA editing enzymes. Taken together, our work expands the scope of programmable RNA editing methods with broad implications for therapeutic modulation of RNA behavior.

## INTRODUCTION

Programmable RNA editing has emerged as a powerful tool for the post-transcriptional manipulation of gene expression^1^. The most widely used systems for targeted RNA editing rely upon adenosine deaminases acting upon RNA (ADARs)^2^, which catalyze adenosine-to-inosine (A-to-I) modifications on double-stranded RNA (dsRNA) and can be programmed to correct disease-causing mutations on mRNA transcripts. In addition, methods have been developed for the selective installation/removal of post-transcriptional modifications, including N^6^-methyladenosine (m^6^A)^3^ and pseudouridine (ψ)^4, 5^, which do not explicitly alter the genetic code but can modulate translation and mRNA metabolism. RNA editing offers a transient approach to gene expression regulation, which can complement genome editing strategies when permanent editing is undesirable^6–8^. Further, the vast repertoire of RNA-modifying enzymes in biology provides a versatile toolbox for tailoring mRNA behavior^9^. Developing efficient and general approaches to selectively manipulate cellular RNA chemistry with enzyme-based platforms has important applications for therapeutic targeting of mRNA^8^.

Current methods for targeted RNA editing/modification offer different solutions to common challenges – delivery of exogenous editing components, recognition of an individual mRNA, and efficient enzymatic transformation at a single site. One class of approaches fuses RNA-modifying enzymes to catalytically inactive CRISPR/Cas-sgRNA complexes (e.g., the RNA-targeting catalytically inactive dCas13), such as the REPAIR/RESCUE methods^10, 11^ developed by Zhang and co-workers that rely upon the human adenosine deaminase ADAR2 and an evolved ADAR2 mutant that performs C-to-U deamination. Delivery of dCas-editor fusion protein and corresponding sgRNA is accomplished using plasmid, mRNA, or viral delivery vector^12^, and site-selective editing is mediated by CRISPR/Cas recognition and formation of a dsRNA with A:C mismatch between sgRNA and target mRNA that directs ADAR activity to a single nucleotide by mimicking its preferred substrate. CRISPR/Cas approaches are general since, in principle, any RNA-modifying enzyme of interest can be fused to dCas13 (or other RNA-targeting CRISPR/Cas protein), and mRNA recognition is readily programmable through the sgRNA sequence; reported dCas13 fusions include m^6^A methyltransferase METTL3^3^, m^6^A demethylases FTO^13^ and ALKBH5^14^, base editor Tad8e^15^, proximity labeling enzymes^16^, translation factors^17^, and others^18^. One challenge to the therapeutic application of dCas13-based RNA editors is delivery, as the large dCas13-fusion protein (typically >100 kDa) and sgRNA must be encoded in a size-constrained gene delivery vehicle^12^, however, efforts towards developing miniaturized Cas proteins that enable packaging of RNA editing machinery into a single adeno-associated virus (AAV) are underway^19, 20^. Further, while mRNA selectivity is programmed by sgRNA sequence, achieving individual site selectivity can still be challenging^12^.

An alternative to CRISPR/Cas-based editing systems is approaches that rely upon chemically modified antisense oligonucleotides (ASOs)^21–29^ or genetically encoded RNA guide strands^30–35^ to direct the activity of an RNA editing enzyme. In contrast to CRISPR/Cas strategies, which are generalizable to a variety of RNA-modifying enzymes, these strategies are largely restricted to directing the A-to-I editing activity of ADAR enzymes. As in the RESCUE/REPAIR strategies described above, ADAR enzyme activity can be programmed by mimicking its native substrate^36, 37^ by the formation of a dsRNA duplex with a mismatched base-pair between the mRNA target and synthetic oligonucleotide or guide RNA. An attractive feature of these approaches for therapeutic application is the ability to redirect the RNA editing activity of endogenous ADAR enzymes, obviating the need for delivery of exogenous RNA editing enzymes or other proteins to cells. Indeed, several RNA editing therapeutics based upon modified ASOs and the activity of endogenous ADAR enzymes are currently in clinical trials^38^, with additional examples likely to enter clinical evaluation soon.

Despite the promise of ASO- and guide RNA-based ADAR editing strategies, generalizing these methods beyond ADAR enzymes is not straightforward, as all methods rely upon the creation of an artificial dsRNA duplex and targeted mismatches to direct deaminase activity. Therefore, it is unknown whether analogous approaches (not relying upon CRISPR/Cas recruitment strategies) can be adapted to reprogram the behavior of other RNA-modifying enzymes, including cytidine deaminases (C-to-U editing) and other post-transcriptional modifiers, which would dramatically expand the therapeutic potential of targeted RNA editing/modification approaches. Further, ADAR-mediated A-to-I editing is limited by the inherent substrate specificity of ADAR enzymes, which restricts efficient editing primarily to UAN base triplets in dsRNA harboring an A:C mismatch, with poorer editing at GAN and CAN triplet motifs^36, 39^.

One approach for targeted RNA editing and modification that enables application beyond ADAR enzymes is the use of self-labeling enzymes to generate chimeric deaminase-ASO fusions in cells. The SNAP-ADAR^21, 27, 29^ method was developed by Stafforst and co-workers and employs a SNAP-tag^40^ fusion of ADAR1/2 (expressed ectopically) together with O^6^-benzylguanine-modified ASOs to mediate efficient and selective RNA editing. Whereas the SNAP-ADAR approach still relies upon the generation of an artificial dsRNA substrate with A:C mismatch to direct ADAR editing, ASO-based recruitment through SNAP-tag (or other self-labeling enzyme chemistry) can direct proximity-based mRNA modification with APOBEC cytidine deaminases^25^, although editing efficiency and programmability for APOBEC-based systems are worse than for ADAR. Application of self-labeling enzymes with ADAR and APOBEC deaminases also requires ectopic expression of the fusion protein, which complicates application of these systems in a therapeutic context and increases off-target editing^29^.

In addition to ADAR and APOBEC family RNA deaminases, directed evolution campaigns starting from the *E. coli* tRNA adenosine deaminase TadA^41^ have yielded a set of highly active and general adenine and cytosine deaminase enzymes for genome editing, known as “base editors”^42, 43^. These enzymes have been principally applied as CRISPR/Cas fusions for targeted genome editing; however, multiple studies have shown that evolved TadA-based deaminases can also deaminate RNA^15, 44–47^. Indeed, proximity-based recruitment of TadA-based editors using RNA-binding protein fusions indicates that these proteins edit RNA substrates with greater efficiency and generality than ADAR and APOBEC enzymes^45, 47^. Further, in contrast to ADAR and APOBEC enzymes, evolved TadA-based editors are compact proteins that can be readily expressed in common heterologous expression systems such as *E. coli*. These features are attractive for therapeutic programmable RNA editing; however, a systematic evaluation of the efficiency and selectivity of TadA-derived deaminases for induced proximity-based RNA editing has not been conducted, and available methods for directing these enzymes to a single mRNA target have so far relied upon CRISPR/Cas strategies^48^.

Herein, we use ASO-based recruitment with the HaloTag^49^ system to evaluate proximity-based RNA editing with evolved TadA-based deaminases. We find that ASO-mediated targeting of TadA-based adenine base editors (ABE) facilitates efficient and localized A-to-I editing on plasmid-based mRNA reporters and endogenous mRNA transcripts. We develop a bulge-forming ASO recruitment architecture that exploits the preference of TadA-based editors for ssRNA substrates and mitigates off-target editing events and can also be applied to APOBEC cytidine deaminases. We also show that off-target editing can be tuned by the selection of evolved TadA-based deaminases with varying activity/promiscuity. Further, we demonstrate that high-level targeted RNA editing is achievable via direct delivery of a recombinant protein-ASO covalent complex formed in the test tube, which decreases off-target transcriptome-wide editing by >99% compared to ectopically expressed editing enzymes that are encoded genetically. Finally, we use our strategy to edit a disease-relevant nonsense mutation in the tumor suppressor PTEN, restoring PTEN expression and highlighting its therapeutic potential. Taken together, ASO-mediated targeting and delivery of TadA-based deaminases provides a versatile and scalable platform for therapeutic RNA editing and modification.

## RESULTS

### ASO-directed RNA editing with TadA8.20

Building upon precedent with SNAP-ADARs^21, 22, 27, 29^, we explored bioconjugation and ASO-based targeting of evolved TadA-derived base editors using the HaloTag system^49^. We synthesized a chloroalkane-modified version of the 18 nt fully 2’-methoxyethyl (MOE)-modified therapeutic ASO Spinraza^50^ (“Halo-Spinraza 1”) with a phosphodiester backbone (rather than the phosphorothioate backbone in the drug) (Fig. 1a). We appended the chloroalkane Halo ligand at the 5’ terminus of the ASO by incorporation of a 5’-amino modifier during solid phase synthesis, followed by reaction with NHS-Halo ligand (Supplementary Information, Supplementary Scheme 1, Supplementary Table 1). Next, we generated Flp-In TRex 293 cells that express a tetracycline-inducible RNA editor-HaloTag fusion (Supplementary Fig. 1). We chose the engineered adenosine deaminase enzyme TadA8.20^44^, which has been shown to possess efficient RNA editing activity^46^. We co-transfected Halo-Spinraza 1 together with a plasmid expressing an mRNA reporter containing the complementary 18 nt Spinraza binding sequence. After 24 h, we observed 41.6% *in cellulo* crosslinking between TadA8.20-HaloTag fusion protein and Halo-Spinraza 1 by Western blot (Fig. 1b). Next, we quantified A-to-I editing^51^ in a 250 bp region centered around the Spinraza binding site by RT-PCR followed by Sanger sequencing. We identified 21 A-to-I editing events demonstrating statistically significant mutation rates as compared to control cells expressing only the TadA8.20-HaloTag fusion (Fig. 1c). The major site of editing, with 68.6% A-to-I conversion, occurred at the first A residue immediately 3’ downstream of the ASO binding site (adjacent to the 5’ end of the ASO where TadA8.20-HaloTag protein is tethered). Statistically significant editing sites were also detected both 5’ upstream of the ASO binding site as well as further downstream in the 3’ direction, but editing efficiency was <25% for these sites, and editing events were considerably reduced in frequency and conversion efficiency at a distance >50 nt away from the ASO binding site.

**Figure 1.**
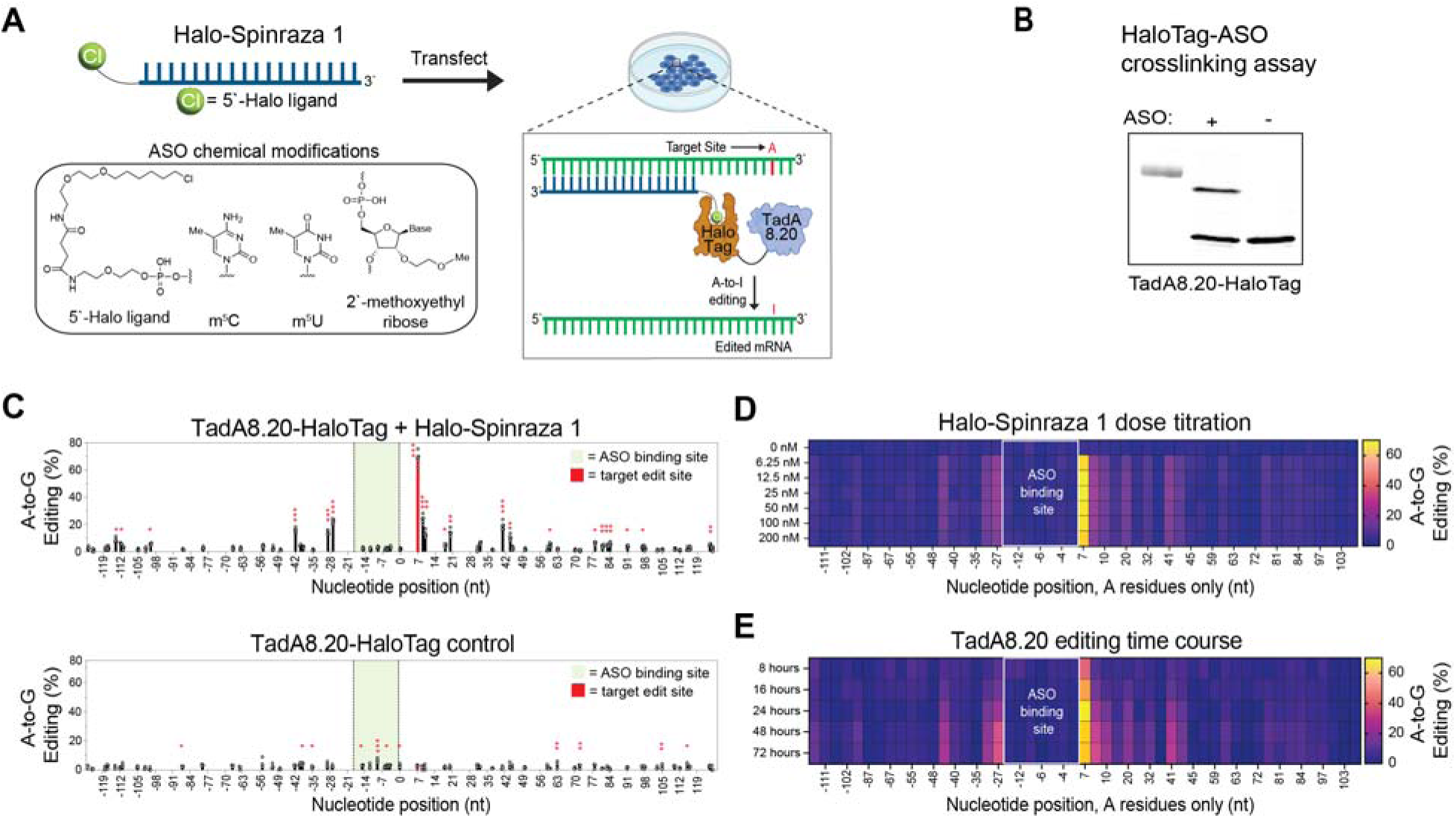
ASO-directed RNA editing with TadA8.20-HaloTag. **(A)** Structure of chloroalkane-modified ASOs and scheme for ASO-directed TadA8.20 editing. **(B)** Western blot analysis of in-cell conjugation between Halo-Spinraza 1 and TadA8.20-HaloTag. **(C)** A-to-G editing of a plasmid-encoded reporter mRNA. TadA8.20-HaloTag-expressing cells were transfected with 25 nM Halo-Spinraza 1 and a plasmid encoding the mRNA reporter sequence. Five independent biological replicates were performed. **(D)** Heatmap showing A-to-G editing on mRNA reporter with different concentrations of Halo-Spinraza 1 in TadA8.20-HaloTag-expressing cells. Data represents the average of two independent biological replicates (Supplementary Fig. 2). **(E)** Heatmap showing time course of A-to-G editing on mRNA reporter sequence with Halo-Spinraza 1 in TadA8.20-HaloTag-expressing cells. Data represents the average of two independent biological replicates (Supplementary Fig. 4). For **(C)-(E),** the x-axis indicates nucleotide position on the mRNA reporter relative to the 5’-end of the Halo-Spinraza 1 binding site, denoted as position 0. The target editing site is colored red. The ASO binding region is shaded green. Editing was quantified from Sanger sequencing data of the mRNA reporter after RT-PCR. Statistical significance legend: * = p ≤ 0.05, ** = p ≤ 0.01, *** = p ≤ 0.001.

We next investigated the effect of ASO concentration on RNA editing. We performed a dose titration with Halo-Spinraza 1 and observed minimal concentration dependence. Editing levels at the major target site immediately 3’ to the ASO binding site were nearly constant from 6.25 nM (63.6% editing) to 200 nM ASO (61.8% editing) (Fig. 1d, Supplementary Fig. 2). Similarly, production of crosslinked ASO-editor complex was largely invariant over this ASO concentration range (Supplementary Fig. 3). Editing levels did decrease marginally at concentrations higher than 25 nM, suggesting a mild Hook effect. Maximal editing was observed at 12.5 nM (68.8%) and 25 nM (68.6%), and 25 nM was used in all further experiments. We also investigated the effect of time on RNA editing. We assessed editing after 8, 16, 24, 48, and 72 h post-transfection (Fig. 1e, Supplementary Fig. 4). After 8 h, editing reached 38.2% at the major edit site, and editing at other positions was low (<10%). Editing increased to 51.9% after 16 h, with editing at secondary edit sites increasing but <15%. We found maximal editing at 24 h (68.9%) and then slowly decreasing editing levels thereafter (61.9% at 48 h; 59.8% at 72 h). Interestingly, in contrast to the major site, the editing of other A sites continued to increase over time. Levels increased from 8.3% at 8 h to 35.0% at 72 h for the second-most highly edited site, which is the first A 5’ upstream of the ASO binding region. Taken together, our data show that ASO-directed proximity-mediated RNA editing can enable selective and high-efficiency RNA editing events with evolved TadA-derived deaminase enzymes.

### ASO-directed RNA editing with a panel of RNA/DNA editing enzymes

We next evaluated the editing behavior of different deaminase enzymes with Halo-Spinraza 1. We selected a panel of adenine and cytosine editors, including the RNA deaminases ADAR2 and APOBEC1, and the evolved DNA base editors TadA8r^52^, TadA7.10^42^, TadDE^53^, and TadCBE^53^. We generated inducible Flp-In TREx 293 cells containing HaloTag-editor fusions and co-transfected Halo-Spinraza 1 together with a plasmid mRNA reporter as described above for TadA8.20. We validated conjugation between Halo-Spinraza 1 and each HaloTag-deaminase fusion protein by Western blot (Supplementary Fig. 5). Editing activity was quantified by Sanger sequencing following RT-PCR.

Among evolved TadA-derived adenine base editors, we observed similar patterns of A-to-I editing (Fig. 2a-2c, Supplementary Fig. 6), with 32.5-51.6% A-to-I editing at the major editing site (3’ to the ASO binding site) and secondary editing <11.9% at all other sites. Whereas editing levels at the major site were lower for these enzymes than for TadA8.20 (Fig. 1c), the prevalence of secondary edits was also reduced, indicating a trade-off between selectivity and activity. To further demonstrate this, we evaluated the effect of time on RNA editing with TadA7.10^42^ a less active precursor to TadA8.20. Unlike TadA8.20, editing at the primary site with TadA7.10 continued to increase until 72 h, achieving 48.8% A-to-I conversion, with editing at other sites <10% (Supplementary Fig. 7). This data also suggests that editor-ASO fusions remain active in cells up to 72 h and shows how editing efficiency and selectivity with TadA-derived adenine editors can be readily tuned by choosing among available enzyme variants.

**Figure 2.**
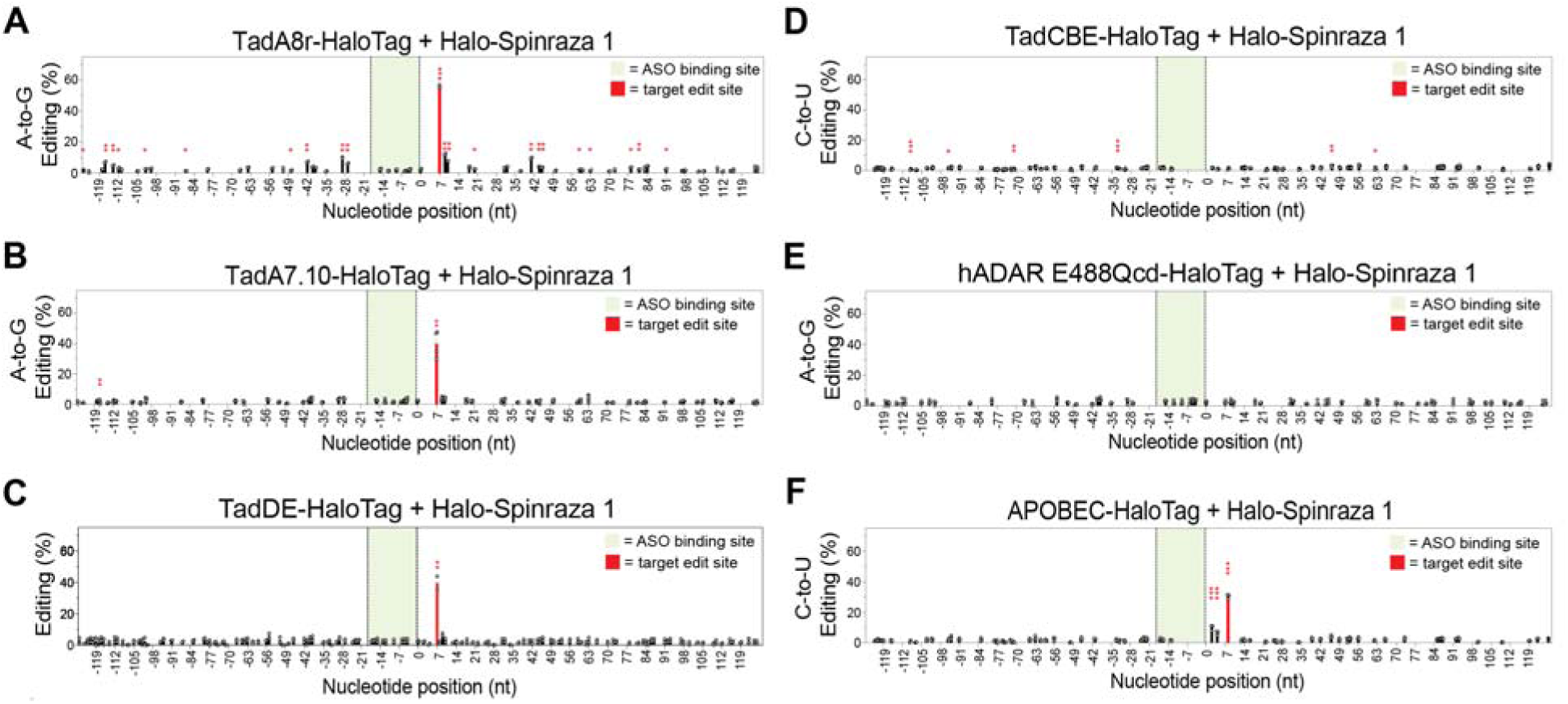
ASO-directed RNA editing with natural and evolved RNA deaminases. Stable cell lines expressing TadA8r-HaloTag **(A)**, TadA7.10-HaloTag **(B)**, TadDE-HaloTag **(C)**, TadCBE-HaloTag **(D)**, hADAR(E488Q)cd-HaloTag **(E)**, or APOBEC1-HaloTag **(F)** were transfected with 25 nM Halo-Spinraza 1 and plasmid-encoded mRNA reporter as in Figure 1. At least two independent biological replicates were performed for all experiments. For **(A)-(E)**, the x-axis indicates nucleotide position relative to the 5’-end of the Halo-Spinraza 1 binding site, denoted as position 0. The target editing site is colored red. The ASO binding region is colored green. Editing was quantified from Sanger sequencing data of the mRNA reporter after RT-PCR. Statistical significance legend: * = p ≤ 0.05, ** = p ≤ 0.01, *** = p ≤ 0.001.

We also explored TadA-derived base editors reported to catalyze C-to-U editing on DNA. For TadDE (Fig. 2c), a dual function editor that can perform A-to-I and C-to-U editing on DNA^53^, we were unable to observe C-to-U editing, whereas A-to-I editing activity was similar to TadA7.10. Similarly, we did not observe C-to-U editing for TadCBE (Fig. 2d), which was specifically evolved for C-to-U editing on DNA^53^, suggesting that DNA C-to-U activity among TadA-derived editors is not readily transferable to RNA. Additionally, we evaluated the catalytic domain from hADAR2 containing the hyperactive E488Q mutation (hADAR E488Qcd)^54^, and rAPOBEC1 (APOBEC), as these enzymes have been used for targeted RNA editing and proximity-based RNA-protein interaction analysis^25, 55, 56^. As expected, we observed no significant editing with hADAR E488Qcd (Fig. 2e, Supplementary Fig. 6e), likely due to its requirement for dsRNA substrates. For APOBEC, we observed 30.9% C-to-U editing immediately 3’ of the ASO binding site (Fig. 2f, Supplementary Fig. 6f), indicating the promise of proximity-directed APOBEC editing, consistent with prior studies^25^. These data demonstrate that whereas localized ASO-directed RNA editing can be achieved with both native APOBEC and evolved TadA-derived enzymes, editing efficiency is considerably higher with TadA-derived deaminases, with minimal compromise in selectivity.

### Preferred editing substrate motif for TadA8.20

We hypothesized that editing selectivity in our system was likely a product of both proximity-mediated ASO recruitment and substrate specificity of the editing enzyme. WT *E. coli* TadA selectively modifies the sequence UACG^41^ within the anticodon loop of tRNA^Arg2^, and evolved TadA-based editors have been reported to preferentially deaminate TA/UA sequence motifs^48, 52, 57^. Within our mRNA reporter plasmid, the major editing site modified by TadA-based editors occurs within the 5mer sequence GTACA, and therefore, we investigated how varying this sequence affects A-to-I conversion. We constructed editing reporter variants containing single, double, or triple mutations (Fig. 3a) and evaluated editing with Halo-Spinraza 1 and TadA8.20-HaloTag. Our data showed a preference for editing at TA dinucleotides, as all three 5mer sequences with this motif (i.e., ATACA, CTACA, and GTATG) were edited between 59.7-69.4% (Fig. 3b-d, Supplementary Fig. 8), similar to the parent GTACA sequence (Fig. 1c). Mutation to GA, AA, or CA (i.e., GGAGG, GAACA, GCACA) (Fig. 3e-g, Supplementary Fig. 8) reduced editing levels to between 22.8-34.6%. Our data suggests that mutation of bases 5’ or 3’ to the TA dinucleotide has minimal impact on editing within this substrate.

**Figure 3.**
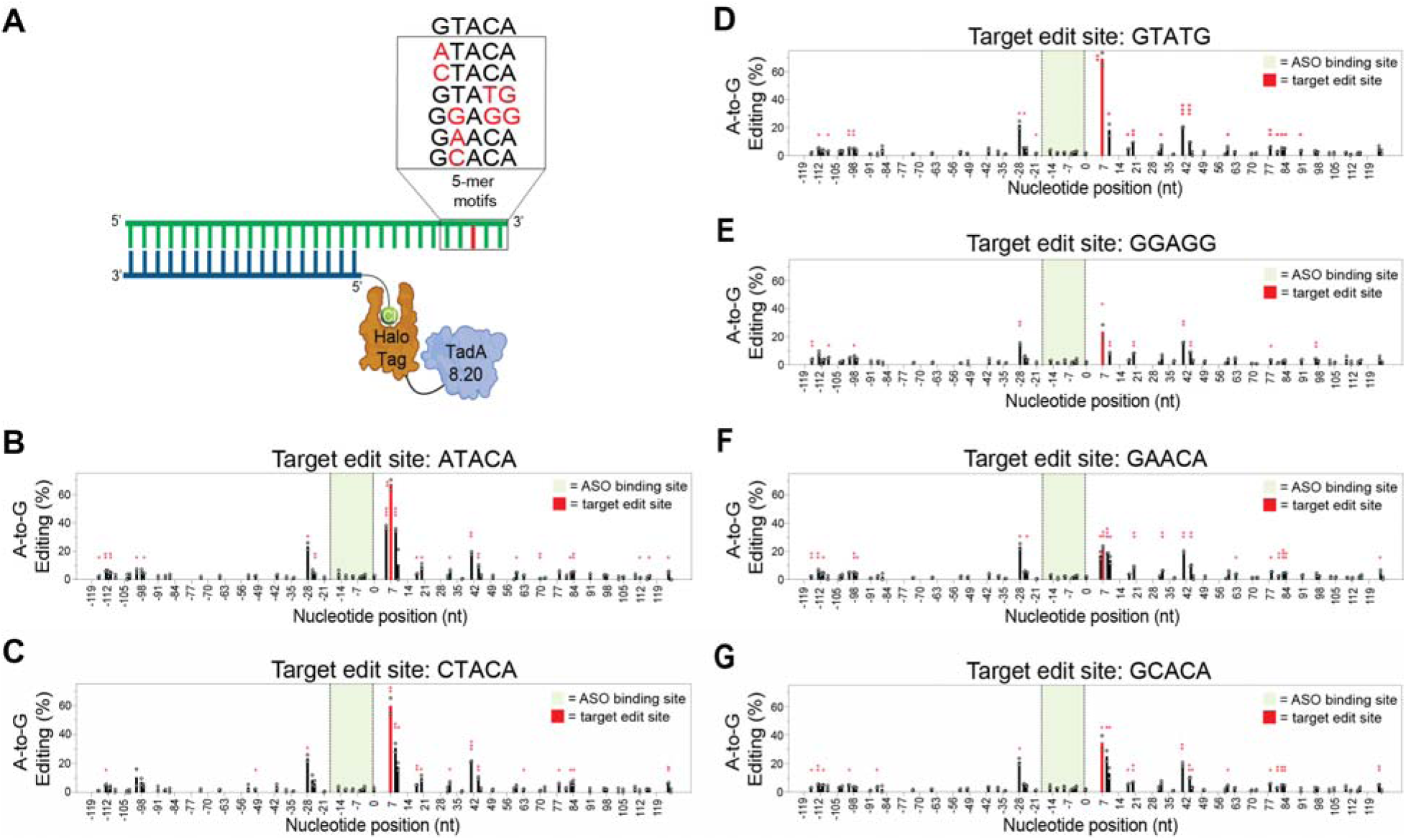
TadA8.20 preferentially edits TA dinucleotides. **(A)** Scheme of assayed sequence motifs. **(B)**-**(G)** Editing of different sequence motifs in TadA8.20-HaloTag-expressing cell line transfected with Halo-Spinraza 1 and plasmid-encoded mRNA reporter. Experiment was performed as in Figure 1. Two independent biological replicates were performed for all experiments. For **(B)-(E),** the x-axis indicates nucleotide position relative to the 5’-end of the Halo-Spinraza 1 binding site, denoted as position 0. The target editing site is colored red. The entire ASO binding region is colored green. Editing was quantified from Sanger sequencing data of the mRNA reporter after RT-PCR. Statistical significance legend: * = p ≤ 0.05, ** = p ≤ 0.01, *** = p ≤ 0.001.

### ASO-directed RNA editing of endogenous mRNA

Having demonstrated editing on a plasmid-based reporter mRNA, we next evaluated editing on endogenous mRNA. We chose ACTB as a model system and synthesized chloroalkane-modified ASOs (ACTB ASO 1-6) (Supplementary Table 1) targeting 6 sites across the coding sequence (CDS) and 3’-UTR. We designed ASOs with similar thermodynamic characteristics to Halo-Spinraza 1 (Supplementary Table 2) and targeted a “TA” motif 7-9 nt downstream of the ASO binding site. We then transfected ACTB ASOs into the TadA8.20-HaloTag-expressing cell line and measured editing using ACTB-specific RT-PCR primers (Supplementary Table 3) and Sanger sequencing, as described above. We observed ASO-dependent editing for all six ASO constructs tested, with the major edit site occurring 3’ of the ASO binding site, consistent with tethering of TadA8.20-HaloTag at the 5’ end of the ASO (Fig. 4, Supplementary Fig. 9). The two ASOs that bind to the 3’-UTR (ACTB ASO 1 and ACTB ASO 2) resulted in 36-56% editing at adjacent A residues (Fig. 4b and 4c), comparable to editing efficiency observed in our plasmid-based reporter mRNA (Fig. 1c). Of these, ACTB ASO 1, which binds a site used in a previous ADAR study^23^, edited multiple A residues 3’ to the ASO binding site at high efficiency (7-56%) (Fig. 4b, Supplementary Fig. 9a). We also observed editing at these sites in the absence of ASO, suggesting they are edited by TadA8.20-HaloTag independent of ASO-directed recruitment (Supplementary Fig. 9a). We were able to mitigate promiscuous editing by applying ACTB ASO 1 in our TadA7.10-HaloTag expressing cell line (Supplementary Fig. 10), however, target site editing efficiency was also modestly decreased to 23.0%. In contrast, for ACTB ASO 2, which also binds in the 3’-UTR, we observed a single major editing site modified at 36.7% (Fig. 4c, Supplementary Fig. 9b) and minimal editing in the absence of ASO. We speculate that the absence of off-target editing with ACTB ASO2 likely stems from a lack of additional substrate sites in this region, but may also depend upon ASO binding ability. Minor editing (<12%) does occur 5’ upstream of the ACTB ASO 2 binding site at other TA dinucleotide motifs (Fig. 4c).

**Figure 4.**
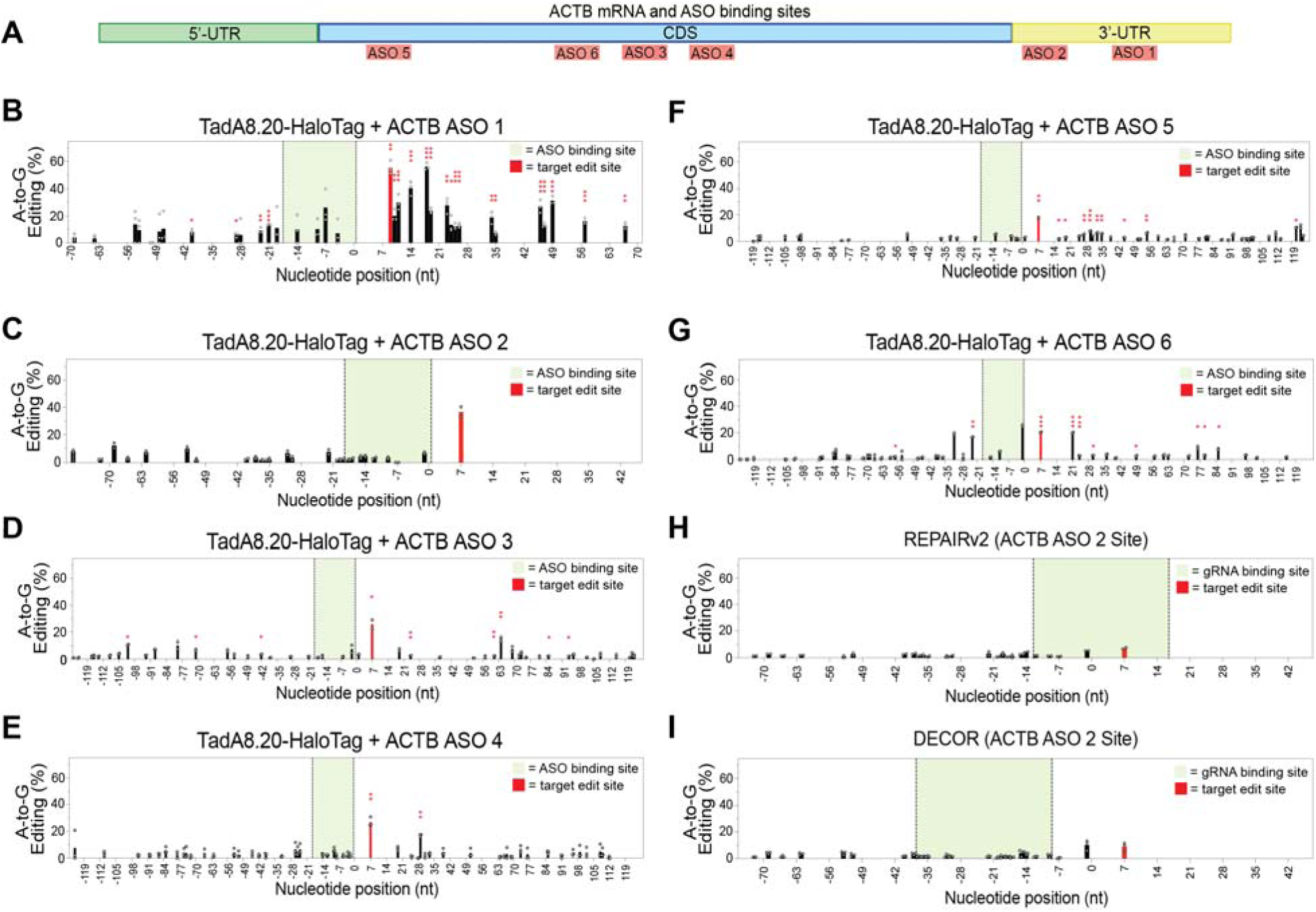
ASO-directed editing on ACTB mRNA. **(A)** Schematic showing the location of ACTB ASO 1-6 binding sites. **(B)**-**(G)** Editing of ACTB mRNA in TadA8.20-HaloTag-expressing cells transfected with ACTB ASO 1-6. **(H)** Editing of the ACTB ASO 2 target site with REPAIRv2 in WT HEK293T cells. **(I)** Editing of the ACTB ASO 2 target site with DECOR in WT HEK293T cells. At least two independent biological replicates were performed for all experiments. For **(B)-(I),** the x-axis indicates the nucleotide position relative to the 5’-end of the ACTB ASO binding site, denoted as position 0. The target editing site is colored red. The ASO **(B)-(G)** or gRNA **(H)-(I)** binding region is colored green. Editing was quantified from Sanger sequencing data of the ACTB mRNA after RT-PCR. Statistical significance legend: * = p ≤ 0.05, ** = p ≤ 0.01, *** = p ≤ 0.001.

ASOs binding the CDS generally showed lower levels of editing than 3’-UTR-binding ASOs, consistent with previous reports^23^. ACTB ASO 3 produced 25.8% editing at an adjacent GTATG sequence with minor editing elsewhere (Fig. 4d, Supplementary Fig. 9c). Similarly, ACTB ASO 4 directed 26.8% editing at an adjacent CTACG sequence with one other major edit (Fig. 4e, Supplementary Fig. 9d). ACTB ASO 5 produced 17.8% editing at the adjacent CTATG site with several low-level but statistically significant editing events downstream (Fig. 4f, Supplementary Fig. 9e). ASO 6 produced 20.4% editing at the adjacent GTACG sequence with one major edit upstream of the ASO binding site (16.7%) and one major edit further downstream (20.1%) along with other low-level edits (Fig. 4g, Supplementary Fig. 9f). In the cases of ACTB ASO 4 and ACTB ASO 6 targets, we observed several A-to-G mutations that were independent of ASO and found in WT cells lacking TadA8.20-HaloTag. These are likely due to the amplification of related actin transcript isoforms (Supplementary Fig. 11). Taken together, our results demonstrate efficient proximity-based ASO-directed editing at multiple sites on an endogenous mRNA and indicate that single-site editing can be achieved through judicious selection of ASO binding site and surrounding sequence.

### Comparison against CRISPR/Cas-directed RNA editing methods

To benchmark our system, we compared it against CRISPR/Cas-based editing strategies, as these methods are compatible with multiple different RNA editing/modifying enzymes, and have been used with TadA-derived base editors for RNA editing^15^. We investigated editing at the ACTB ASO 2 target site using REPAIRv2^10^, an ADAR method, and DECOR^15^, which uses TadA8e (a TadA-derived adenine base editor with activity intermediate between TadA7.10 and TadA8.20)^44^. We chose CRISPR sgRNAs for each method following literature precedent. For REPAIRv2, the sgRNA contained an A:C mismatch over the target site positioned 20 nt from the 3’-end of the sgRNA; using this construct we observed only 7.4% A-to-I editing, 5-fold lower than ASO-directed editing with TadA8.20 (Fig. 4c and 4h, Supplementary Fig. 12a). Using the DECOR method, we designed a CRISPR sgRNA positioning the target editing site 15 nt from the 5’-end of the sgRNA and also observed poor editing (9.0%) (Fig. 4i, Supplementary Fig. 12b). Whereas REPAIRv2 and DECOR have been validated on other mRNA substrates, our results show that these methods are context-dependent (or require empirical optimization of sgRNA sequence), and highlights an mRNA transcript context where our ASO-guided approach performs more efficiently. The DECOR system also features dual SV40 nuclear localization sequences (NLS), reported to result in higher mRNA editing^15, 58^. We tested an analogous TadA8.20-NLS-HaloTag-NLS construct via transient transfection with ACTB ASO 2 and measured 27.4% editing, slightly lower than editing without the dual NLS sequences (Supplementary Fig. 13 and Fig. 4c).

### RNA editing via cellular delivery of a recombinant deaminase-ASO complex

To extend the application of our method to cells that do not require genetic engineering or transgene delivery and reduce off-target editing, we hypothesized that cellular delivery of a recombinant TadA8.20-ASO oligo-protein complex formed in the test tube could mediate site-specific RNA editing (Fig. 5a), as an analogous strategy has been successful for delivering Cas9-sgRNA RNA-protein (RNP) complexes for genome editing^59–62^. Based on precedent with Cas9-sgRNA RNPs^59–66^, we explored TadA8.20-ASO delivery by electroporation or liposome transfection. We first generated recombinant Tad8.20-HaloTag protein through heterologous expression in *E. coli* and demonstrated efficient *in vitro* labeling with a choroalkane-modified ASO (Supplementary Figs. 14 and 15). We next formed TadA8.20-HaloTag-ASO conjugates with ACTB ASO 1, 2, or 3, as these ASOs performed well in the Tad8.20-HaloTag-expressing cell line (Fig. 4) and delivered the oligo-protein conjugates into WT HEK293T cells.

**Figure 5.**
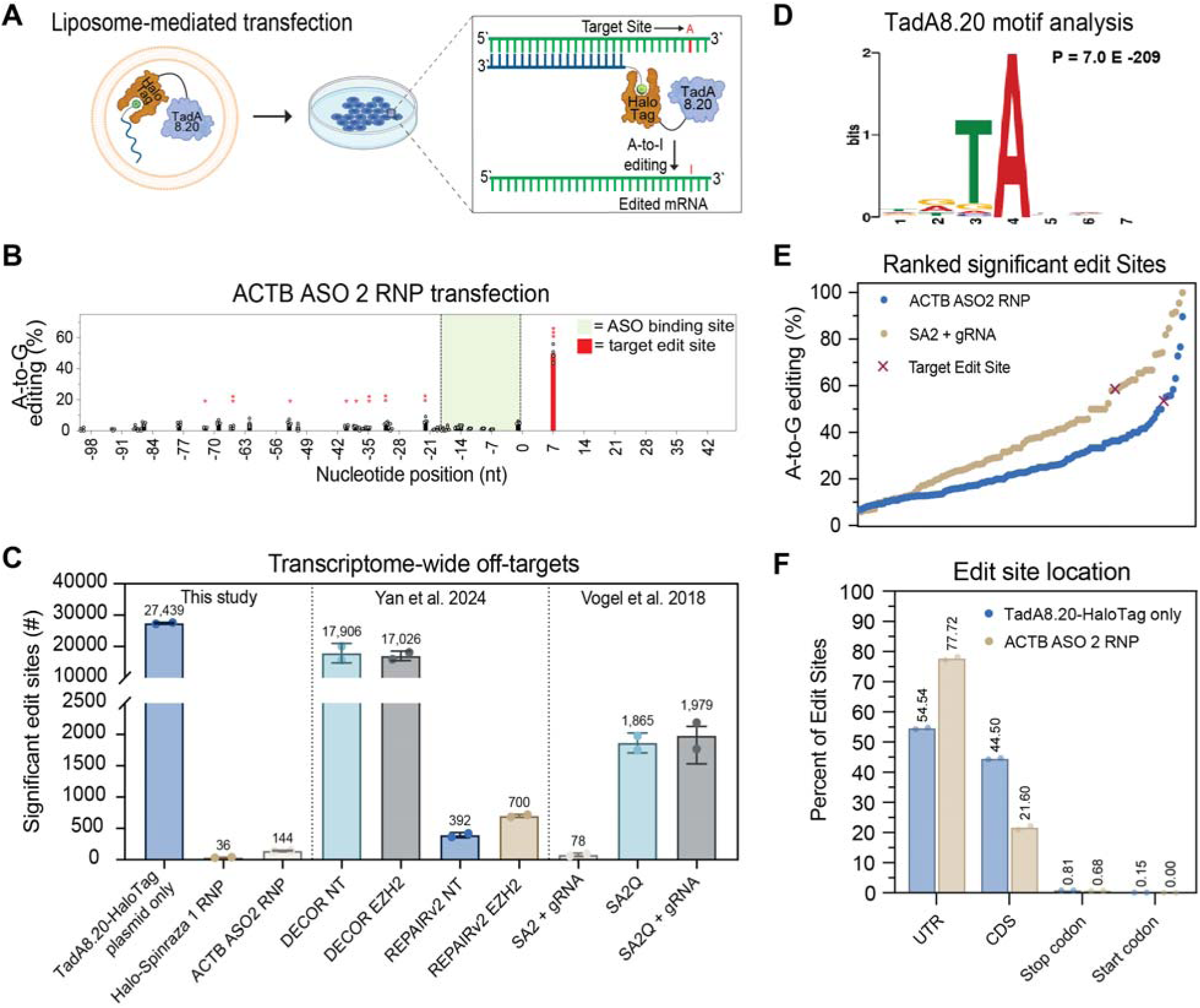
RNP delivery reduces transcriptome-wide off-target editing. **(A)** Schematic illustrating RNP complex LNP delivery and targeted editing in unmodified cells. **(B)** Editing on ACTB mRNA with ACTB ASO 2 RNP complex transfected into HEK293T cells using Lipofectamine 2000. The x-axis indicates the nucleotide position relative to the 5’-end of the ASO binding site, denoted as position 0. The target editing site is colored red. The ASO binding region is colored green. Editing was quantified from Sanger sequencing data of the ACTB mRNA after RT-PCR. Four independent biological replicates were performed. **(C)** Transcriptome-wide off-target edit sites detected in TadA8.20-HaloTag plasmid only, Halo-Spinraza 1 RNP, and ACTB ASO 2 RNP, compared to DECOR, REPAIRv2, and SNAP-ADAR. DECOR and REPAIRv2 sequencing data was obtained from Yan and Tang 2024^15^. SNAP-ADAR sequencing data was obtained from Vogel et al. 2018^29^. w. **(D)** MEME analysis of the preferred substrate motif for TadA8.20-HaloTag. Statistically significant edit sites identified in cells transfected with TadA8.20-HaloTag plasmid were used. **(E)** Histogram of transcriptome-wide editing site conversion in detected sites from ACTB ASO 2 RNP and SNAP-ADAR (SA2). **(F)** Distribution of edit sites based on transcript features for significant detected sites in TadA8.20-HaloTag plasmid and ACTB ASO 2 RNP conditions. For whole-transcriptome RNA-seq, two independent biological replicates were performed and analyzed. Statistical significance legend: * = p ≤ 0.05, ** = p ≤ 0.01, *** = p ≤ 0.001.

We tested different concentrations (25-100 nM) of TadA8.20-HaloTag-ACTB ASO 1 complex (“RNP complex”) with different commercial transfection reagents and electroporation and found similar editing patterns on ACTB mRNA as in our previous experiments with TadA8.20-HaloTag expressing cells transfected with ACTB ASO 1. Editing at the target position measured 24.0% (Supplementary Fig. 16a), about half of what we observed in TadA8.20-HaloTag-expressing cells. We did not observe major differences in editing across the transfection conditions assayed (Supplementary Figs. 16 and 17), and further experiments were performed using Lipofectamine 2000 and 25 nM deaminase-ASO complex. For the ACTB ASO 2 conjugate, we measured 49.7% editing at the primary editing site (Fig. 5b) with minimal secondary editing; the efficiency and specificity, in this case, were better than in the TadA8.20-HaloTag-expressing cell line. Targeting the CDS with the TadA8.20-HaloTag-ACTB ASO 3 complex yielded 8.3% editing at the primary site (Supplementary Figs. 16f-g), lower than editing when the ASO was transfected into TadA8.20-HaloTag-expressing cells (Fig. 4d).

We also evaluated the efficacy of RNP delivery and editing in HeLa cells to establish generality. We observed targeted editing with both ACTB ASO 1 RNP complex and ACTB ASO 2 RNP complex, with 17.8% and 24.8% A-to-I conversion, respectively, at the target position (Supplementary Fig. 18a-d). For both ACTB RNP complexes, editing efficiency was lower in HeLa cells than in HEK293T, which may be attributable to lower transfection efficiency in HeLa cells. Collectively, these data demonstrate editing of endogenous mRNA by direct cellular delivery of a recombinant deaminase-ASO conjugate.

### RNP delivery reduces transcriptome-wide off-target editing

We next investigated whether RNP delivery reduced transcriptome-wide off-target editing as compared to a genetically encoded TadA8.20-HaloTag construct. We prepared RNA-seq libraries for Illumina sequencing from WT HEK293T cells treated with the ACTB ASO 2 RNP complex and analyzed A-to-G mutations (comparing against untreated cells) following literature precedent^15^. We also analyzed A-to-I editing in cells transfected with a plasmid encoding TadA8.20-HaloTag and cells treated with a Halo-Spinraza 1 RNP complex (as a non-targeting control). For Halo-Spinraza 1 RNP transfection, we did not detect A-to-I edits on SMN2 mRNA; however, the transcriptome-wide analysis contained no reads mapped to the intronic region in SMN2 pre-mRNA where Spinraza binds, likely due to the short lifetime of the pre-mRNA species and the polyA-enrichment step used to prepare mRNA for RNA-seq library preparation.

We found that RNP delivery resulted in ∼100-1,000-fold reduction in transcriptome-wide off-target editing events as compared to plasmid-based expression of TadA8.20-HaloTag. Transfection of cells with plasmid-encoded TadA8.20-HaloTag resulted in an average of 27,439 transcriptome-wide A-to-I edits between two independent biological replicates (Fig. 5c, Supplementary Fig. 19a, Supplementary Datafile 1). Although we did not co-transfect a chloroalkane-containing targeting ASO in this condition, we estimate that only 40% of the HaloTag-editor is covalently conjugated to Halo-ASO in cells (Fig. 1b). In contrast, lipofection of Halo-Spinraza 1 RNP and ACTB ASO 2 RNP resulted in 36 and 144 transcriptome-wide editing events, respectively (Fig. 5c, Supplementary Fig. 19a, Supplementary Datafile 1). We also performed comparative analysis of transcriptome-wide off-target editing using published RNA-seq data from the RNA editing methods DECOR, REPAIRv2, and SNAP-ADAR^15, 29^. The transcriptome-wide off-target profiles for ACTB ASO 2 RNP and Halo-Spinraza 1 RNP are most similar to SNAP-ADAR2 (SA2) (78 edit sites) and superior to DECOR, SNAP-ADAR2(E488Q) (SA2Q), and REPAIRv2, which contained 17,036, 1,979, and 673 edit sites, respectively (Fig 5c, Supplementary Datafile 1). Importantly, the major ACTB ASO 2 editing site in the 3’-UTR of ACTB observed by RT-PCR and Sanger sequencing (Fig. 5b) was comparably edited at 49.7% in the transcriptome-wide analysis (Supplementary Fig. 19b). We determined the substrate motif for TadA8.20 by analyzing 27,439 edit sites observed in the TadA8.20-HaloTag plasmid transfection condition, which revealed a preferred “TA” dinucleotide motif (Fig. 5d, Supplementary Datafile 1), corroborating previous findings for the substrate preference of TadA variants^41, 42, 67, 68^ and our findings with different substrate motifs (Fig. 3). We further compared the editing stoichiometry of sites detected in the ACTB ASO 2 RNP condition with those detected in SA2 (Fig. 5e, Supplementary Datafile 1), as SA2 gave a similar number of transcriptome-wide editing events (Fig. 5c). Although the number of off-target sites is similar, the sites detected in SA2 are modified at higher stoichiometry compared to those in ACTB ASO 2 RNP (Fig. 5e, Supplementary Fig. 19c). Editing stoichiometry for ACTB ASO 2 RNP was also lower across most sites compared to TadA8.20-HaloTag (Supplementary Fig. 19c). We also evaluated the transcript location distribution of edit sites detected in ACTB ASO 2 RNP and TadA8.20-HaloTag plasmid conditions. The majority of edit sites occurred in UTRs and CDSs, with a minor fraction occurring in stop and start codons (Fig. 5f). Additionally, ACTB ASO 2 RNP gave more sites that occurred in UTRs and fewer in CDSs relative to TadA8.20-HaloTag (Fig. 5f). Altogether, these data highlight the potential of RNP delivery with covalent deaminase-ASO conjugates as a strategy to achieve efficient, site-specific RNA editing in unmodified cells without off-target transcriptome-wide editing.

### Alternative ASO architectures increase RNA editing selectivity

Whereas RNP delivery dramatically reduced transcriptome-wide off-target editing, transcript-specific secondary editing was largely unaffected (Supplementary Fig. 16). We, therefore, explored alternative ASO architectures to further restrict TadA8.20 activity to the target site of interest. Since mRNA editing was biased towards the 5’-end of the ASO (where the editor is tethered), we evaluated tethering the editing enzyme to a nucleotide located in the middle of the ASO (Fig. 6a). We synthesized a 2’-MOE-modified phosphodiester ASO with the same nucleotide sequence as Halo-Spinraza 1 but installed the chloroalkane Halo ligand at the C5 position of an internal dT residue (Fig. 6a) located 10 nt from the 5’-end of the ASO (“Halo-Spinraza 2”). We evaluated editing on the same mRNA reporter used previously (Figs. 1-3) and observed 49.4% editing at the major target position (as compared to 68.6% with the 5’-Halo-modified Spinraza ASO) but with considerably lower secondary editing at other positions on the transcript (Fig. 6b and Fig. 1c). Building on the improvements conferred by this simple modification in ASO-enzyme conjugate design, we tested a “bulge-forming” ASO architecture in which ASO-mRNA binding is predicted to generate a ssRNA “bulge” opposite the editing enzyme tethered at the middle of the ASO. Since our data and other published reports^15, 41, 67, 69, 70^ indicate that TadA-derived enzymes preferentially modify ssRNA, we reasoned that editing would be biased towards the ssRNA bulge, whereas adjacent bases would be protected by base pairing with the ASO. We designed ASOs with discontinuous binding regions intended to position the target mRNA base within a ssRNA loop varying between 4 to 9 nt, and with the Halo ligand at the middle of the ASO sequence (“Bulge ASO 1-3”) (Fig. 6a). Gratifyingly, editing using the bulge-forming ASOs was efficient and specific for the target base, with a similar profile of on-target to off-target activity observed for the 6 nt and 9 nt bulge architectures as observed for a continuous ASO design with the editor linked at the middle (Fig. 6c-e).

**Figure 6.**
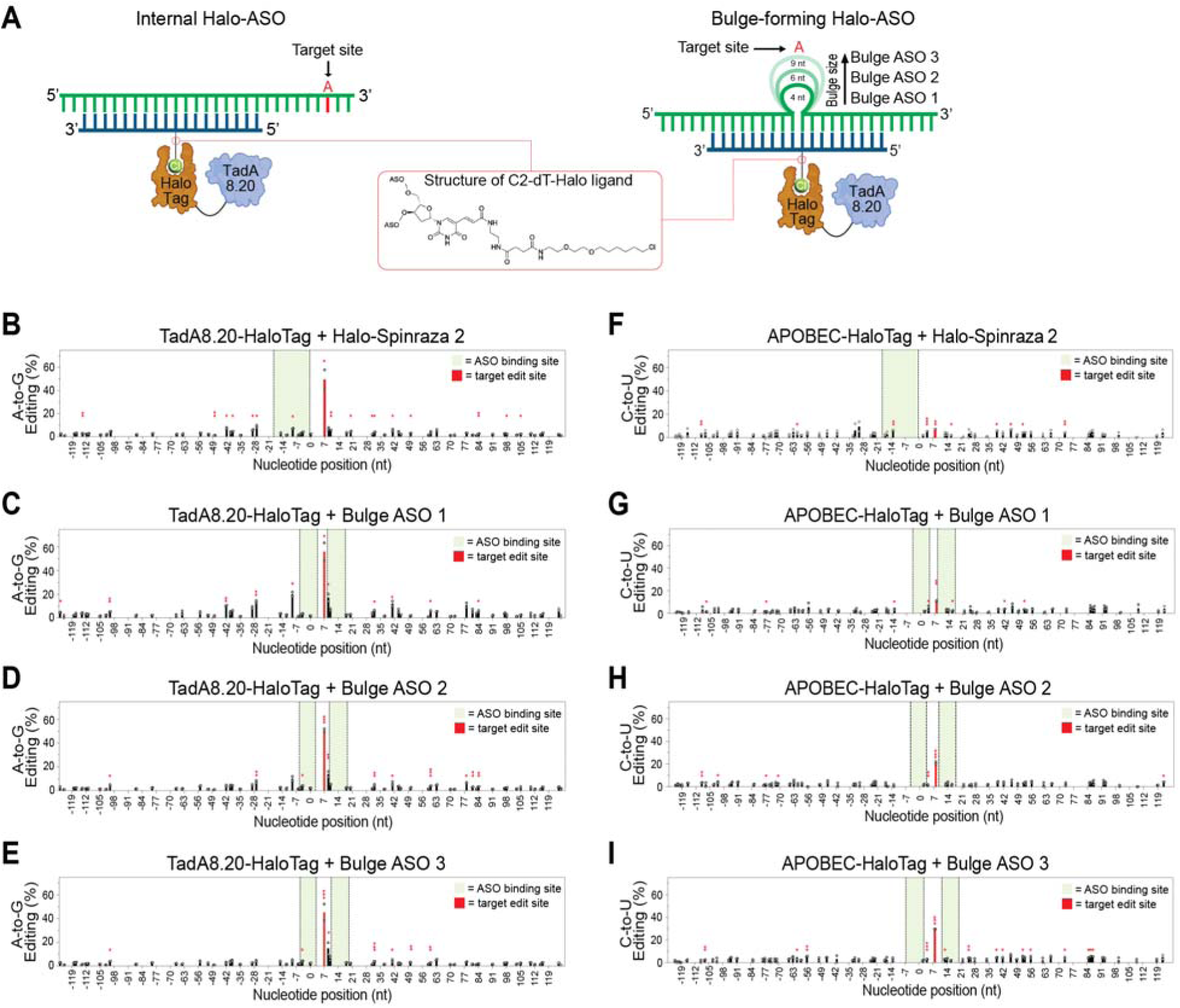
Alternative ASO architectures increase RNA editing selectivity. **(A)** Schematic illustrating alternative ASO designs: an internally modified ASO where the target site remains downstream of the binding site (left), and a bulge-forming ASO where the target site is positioned within a single-stranded loop (right). The structure of the internal dT-Halo ligand modification is boxed in red. **(B)** Halo-Spinraza 2-directed editing of the reporter mRNA transcript in TadA8.20-HaloTag expressing cells. **(C)-(E)** Bulge ASO 1-**(C)**, 2-**(D)**, and 3-**(E)** directed editing of the reporter mRNA transcript in TadA8.20-HaloTag expressing cells. **(F)** Halo-Spinraza 2-directed editing of the reporter mRNA transcript in APOBEC-HaloTag expressing cells. **(G)-(I)** Bulge ASO 1-**(G)**, 2-**(H)**, and 3-**(I)** directed editing of the reporter mRNA transcript in APOBEC-HaloTag expressing cells. For **(B)-(I)**, experiments were performed as in Figure 1. The x-axis indicates nucleotide position relative to the 5’-end of the Halo-Spinraza 2 binding site, denoted as position 0. The target editing site is colored red. The ASO binding region is colored green. Editing was quantified from Sanger sequencing data of the mRNA reporter after RT-PCR. Two independent biological replicates were performed for all experiments. Statistical significance legend: * = p ≤ 0.05, ** = p ≤ 0.01, *** = p ≤ 0.001.

We further evaluated whether alternative ASO architectures were compatible with other RNA deaminases in addition to TadA8.20. For TadA7.10, we measured 16.6% editing (Supplementary Fig. 20a) with Halo-Spinraza 2 at the target position with only one low-level statistically significant secondary edit. Editing with Bulge ASO 2 resulted in 30.2% A-to-I conversion at the target site, also with only one low-level statistically significant secondary edit (Supplementary Fig. 20b), similar to results with the 5’-Halo-ligand-modified Halo-Spinraza 1 (Fig. 2b, 39.8% editing at the target site). Thus, TadA7.10 editing is compatible with internal ASO conjugation and the bulge-forming ASO architecture; however, TadA7.10 demonstrated high target site selectivity with the original 5’-Halo ASO design, and neither ASO design strategy further improved editing efficiency or selectivity in the reporter system

For APOBEC1, a C-to-U deaminase, we found that editing at the target base directed by Halo-Spinraza 2 was low efficiency (7.4%) (Fig. 6f) as compared to Halo-Spinraza 1 (Fig. 2f, 30.9% conversion). In contrast, we found efficient APOBEC1-mediated C-to-U editing directed by bulge-forming ASOs with fewer secondary editing sites. In particular, Bulge ASO 2 and Bulge ASO 3, which induce 6 nt and 9 nt bulges in the mRNA substrate, respectively, resulted in 21-30% editing at the target site (Fig. 6h-i). Editing with Bulge ASO 1, which generates a smaller 4 nt bulge, was less efficient with 10% conversion (Fig. 6g). Importantly, the two secondary edits observed with Halo-Spinraza 1 (Fig. 2f) were eliminated with the bulge-forming ASOs while retaining on-target efficiency. Collectively, these data indicate that off-target editing activity of diverse A-to-I and C-to-U RNA deaminases that preferentially modify ssRNA can be controlled through rational ASO design and modification without compromising on-target editing at the primary site.

### Correction of a disease-causing mutation with ASO-directed RNA editing

Finally, we tested whether our method could correct nonsense mutations in a disease-associated mRNA and restore protein expression (Fig. 7a). Inactivating mutations in PTEN are common in tumors and other germline diseases such as Cowden’s syndrome^71^. We selected the Q245X mutation in PTEN, which yields a truncated and inactive version of the tumor suppressor, as a candidate mutation to test in our system. PTEN Q245X has been subjected to small-molecule drug readthrough experiments with inconsistent results ranging from <1% to 11.7% restoration^72, 73^. We cloned the mutant PTEN gene into a plasmid and co-transfected it into TadA8.20-HaloTag-expressing cells together with either bulge-forming ASOs or a 5’-Halo-modified ASO targeting the mutation site. Editing was measured 24 h after transfection by RT-PCR and Sanger sequencing. The 5’-Halo ASO design (PTEN ASO 1) gave only 9.5% editing (Fig. 7b, 7h, Supplementary Fig. 21a). Using a 7 nt bulge-forming ASO (PTEN ASO 2), we achieved 33.4% editing at the target site (Fig. 7c, 7h), with up to 12.9% editing at other sites within the gene. PTEN ASO 2 with TadA7.10-HaloTag gave only 13.4% editing but eliminated most other statistically significant edit sites (Supplementary Fig. 21b, 21c). Increasing the length of the ASO binding regions in PTEN ASO 2 from 22 nt to 26 nt generated PTEN ASO 3, which increased editing to 36.4% at the target position and up to 18.2% at secondary sites (Fig. 7d, 7h). An ASO construct that reduces the bulge size from 7 nt to 5 nt (PTEN ASO 4) modestly decreased editing efficiency to 23.4% (Fig. 7e, 7h). Importantly, for all PTEN ASOs, the two highest secondary edits at nucleotide positions 30 and 38 relative to the PTEN ASO 1 binding site are conservative mutations, yielding Ile to Val and Val to Val, respectively. These editing approaches rescued PTEN expression by as much as 23.5% of the wild-type control (after 24 h) as evaluated by Western blot (Supplementary Fig. 22).

**Figure 7.**
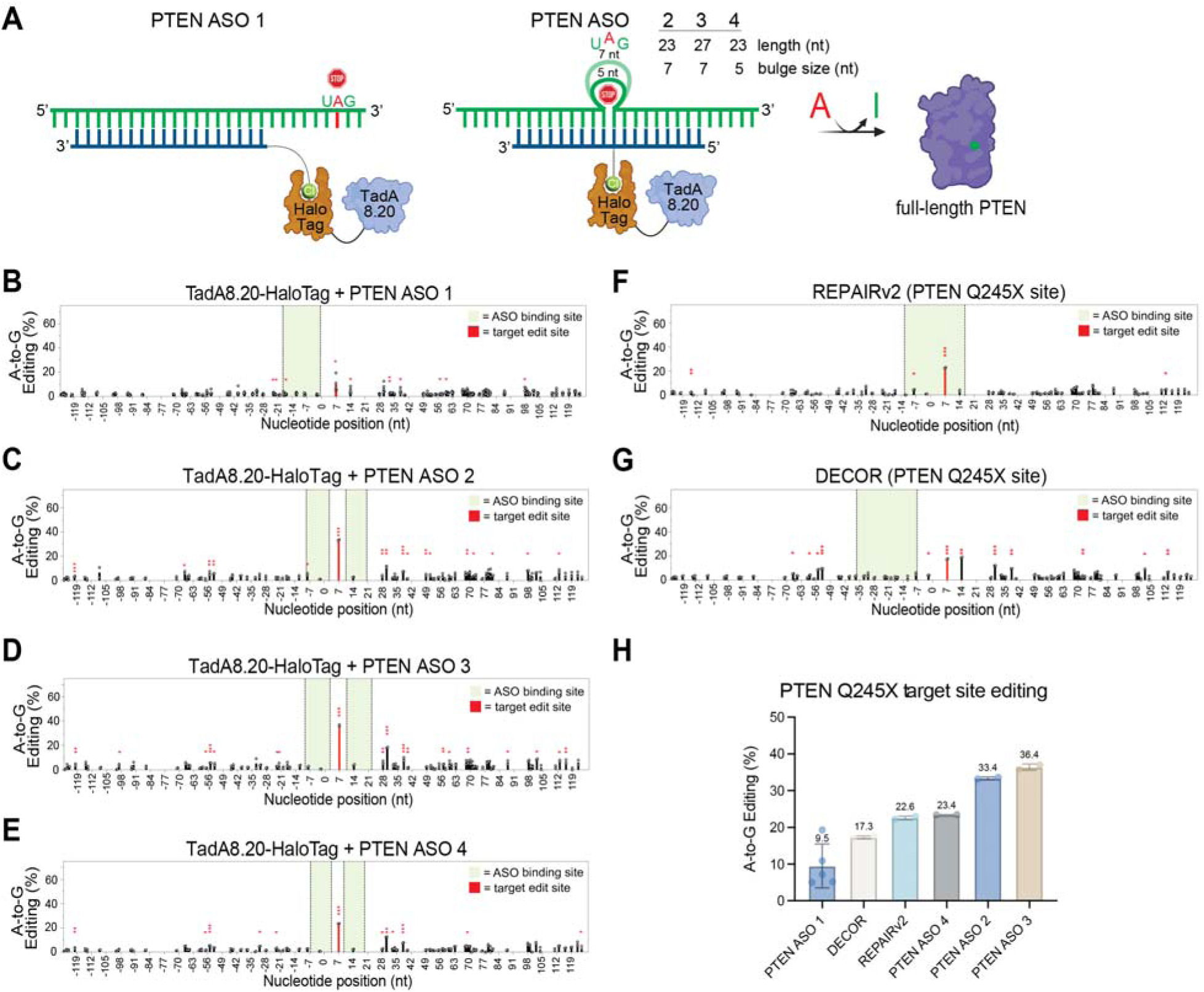
Restorative editing of premature termination codon Q245X in PTEN. **(A)** Schematic illustrating the application of different ASO architectures to edit the premature termination codon and restore protein expression. **(B)** Editing with PTEN ASO 1 in TadA8.20-HaloTag expressing cells. Cells were transfected with PTEN ASO1 and a plasmid encoding PTEN Q245X mutant. **(C)** Editing with PTEN ASO 2 in TadA8.20-HaloTag expressing cells. **(D)** Editing with PTEN ASO 3 in TadA8.20-HaloTag expressing cells. **(E)** Editing with PTEN ASO 4 in TadA8.20-HaloTag expressing cells. **(F)** Editing with REPAIRv2 in WT HEK293T cells. **(G)** Editing with DECOR in WT HEK293T cells. **(H)** Editing percentage at the PTEN Q245X target site using different editing strategies. At least two independent biological replicates were performed for all experiments. For **(B)-(G)**, the x-axis indicates nucleotide position relative to the 5’-end of the PTEN ASO 1 binding site, denoted as position 0. The target editing site is colored red. The ASO **(B)-(E)** or gRNA **(F)-(G)** binding region is colored green. Editing was quantified from Sanger sequencing data of the PTEN Q245X mRNA after RT-PCR. Statistical significance legend: * = p ≤ 0.05, ** = p ≤ 0.01, *** = p ≤ 0.001.

We also evaluated the abilities of REPAIRv2 and DECOR to edit PTEN Q245X. In our hands, both methods were less efficient than our approach at primary site editing, with varying degrees of off-target editing. REPAIRv2 gave 22.6% on-target editing with minimal off-target editing at other positions within the gene (Fig. 7f, 7h, Supplementary Fig. 21d). DECOR gave 17.3% on-target editing with multiple secondary edit sites (Fig. 7g, 7h, Supplementary Fig. 21e). Therefore, our approach performs favorably as compared to CRISPR/Cas methods, with superior on-target editing efficiency.

## DISCUSSION

Here we demonstrate programmable ASO-directed RNA editing with TadA-derived deaminase enzymes. We use the HaloTag system and 2’-MOE-modified ASOs to recruit these enzymes and mediate proximity-directed editing on mRNA substrates of interest, including endogenous and disease-relevant transcripts. Further, we show that a covalent TadA8.20-ASO RNP complex can be delivered directly to cells for targeted RNA editing with minimal transcriptome-wide off-target editing. TadA-derived “base editors” have been generated through directed evolution for genome editing, but our study and others^74–76^ show that these proteins, which are derived from a bacterial tRNA deaminase^41^, have largely retained their RNA deaminase activity and constitute a promising pool of enzymes for RNA editing applications.

In contrast to ADAR proteins, which have been the focus of programmable RNA editing efforts^1^, TadA enzymes recognize ssRNA substrates. Whereas this property can be leveraged to address a wider range of RNA substrates, it also requires new approaches for constraining proximity-directed editing on target transcripts to individual nucleotides. One strategy is to harness the sequence-specificity of TadA proteins – we show herein that editing is preferred on TA dinucleotides. Therefore, an ideal target edit site will contain an isolated TA motif. Another approach is to protect potential secondary edit sites through ASO-mRNA base pairing, as we find that TadA enzymes do not modify RNA bases in the ASO binding site. Towards this end, we develop a new bulge-forming ASO recruitment architecture for TadA proteins that tethers the enzyme in the middle of the ASO and induces an mRNA loop surrounding the target edit site and opposite the conjugated enzyme upon ASO-mRNA binding. Our work shows that this design can reduce secondary editing without compromising editing efficiency as compared to a more conventional ASO design that tethers the editing enzyme to the end of the ASO. Indeed, in the case of programmable editing on a PTEN mutant transcript, the bulge-forming ASO dramatically increases editing efficiency, suggesting that this ASO design may interfere with secondary structure within the transcript that otherwise impedes editing on the target Q245X mutation site. Since tethering the editing enzyme in the middle of the ASO restricts secondary editing as compared to conjugation at the end of the sequence, we propose that systematic evaluation and optimization of the linker chemistry between enzyme and ASO (currently formed by HaloTag) may serve to further constrain editing activity at the target site and increase selectivity. In addition, changing the identity of the RNA editing enzyme in our system is a viable approach to tuning editing efficiency, selectivity, and the nature of the editing mutation. Interestingly, the bulge-forming architecture is amenable to directing the activity of APOBEC C-to-U deaminases, which opens new opportunities for deploying these enzymes in RNA editing systems. In addition, TadA proteins are amenable to directed evolution to generate new derivatives with tunable editing behavior, and a large number of TadA-derived enzymes have been reported in the literature^44, 53, 69, 70, 77–79^. In this work, we rely primarily upon TadA8.20, which is among the most active of reported TadA proteins, but we also demonstrate ASO-directed editing with TadA7.10, which offers increased selectivity at the expense of reduced editing efficiency. Moving forward, the evolution of TadA-based editors specifically for RNA editing^70^, rather than repurposing constructs developed for genomic editing, is likely to provide tailor-made RNA editors that can address the limitations of the current enzymes.

RNA editing with exogenous deaminase enzymes, such as TadA-derived enzymes that are of bacterial origin, necessitates a viable cellular delivery strategy to introduce the RNA editing enzyme and associated targeting constructs. This challenge also exists for CRISPR/Cas-based RNA editing methods such as REPAIR and self-labeling enzyme approaches like the SNAP-ADAR method, and is commonly addressed through transgene delivery on viral vectors (i.e., AAV) or through mRNA/plasmid transfection as lipid nanoparticles (LNPs). Generally, RNA editing approaches that require cellular expression of an exogenous RNA editing enzyme are more prone to off-target transcriptome-wide editing than those that can rely upon endogenous editing enzymes^1, 31, 80^. Inspired by CRISPR/Cas9 genome editing methods that deliver the Cas9-sgRNA complex as an RNP^60–66^, we show that an analogous TadA8.20-ASO complex can be delivered as an LNP, and mediate efficient RNA editing in cells. Not only does this solve the transgene delivery problem without requiring genetic modification of the recipient cells, but we find that transcriptome-wide edits are dramatically reduced using this approach, likely due to lower cellular concentrations of the editing enzyme and stoichiometric formation of the enzyme-ASO complex. We propose that delivery of exogenous RNA editing enzymes with targeting oligonucleotides in the form of covalent RNP complexes may present a general strategy for programmable RNA editing in cells with a variety of editors and the ability to readily titrate dosage. Systematic evaluation of protein, linker, and ASO properties that are compatible with LNP delivery and endosomal escape will provide important insights into the generality of this approach and open the door to the delivery of diverse RNA effector proteins in the form of protein-ASO conjugates. Such studies are underway in our group and will be reported in due course.

## METHODS

### General cell culture

All cells were grown at 37 °C in a humidified atmosphere with 5% CO2 in DMEM (Life Technologies) supplemented with 10% fetal bovine serum (Atlanta), 1x penicillin-streptomycin (Gibco Life Technologies), and 2 mM L-glutamine (Life Technologies).

### Stable cell line generation

To generate stable cell lines expressing editor-HaloTag fusion proteins, Flp-In TRex 293 cells were seeded at 6 × 10^5^ cells per well in six-well plates, and co-transfected with the appropriate pCDNA5/FRT/TO-editor-HaloTag plasmid (0.2 µg) and pOG44 plasmid (1.8 µg, Thermo Fisher). After selection with 100 µg/mL hygromycin B and 15 µg/mL blasticidin, colonies were expanded.

### Plasmid construction

Gene blocks for human codon-optimized deaminases, HaloTag, and PTEN Q245X were obtained from Twist Biosciences, restriction enzyme-digested, and ligated into pcDNA5 vector with T4 DNA ligase (New England Biolabs). Wild-type PTEN control was synthesized by site-directed mutagenesis of the PTEN Q245X and overlap extension PCR. For REPAIRv2 and DECOR gRNAs, vectors for PspCas13b (REPAIR) (Addgene #103854) and RfxCas13d (DECOR) (Addgene #109053) were digested with BbsI-HF (New England Biolabs) and purified following agarose gel electrophoresis. Targeting gRNA oligos were obtained from Sigma, annealed and phosphorylated with T4 PNK (New England Biolabs), diluted 1:200, and 1 μL was ligated into 50 ng vector. For REPAIRv2 gRNA design, the A:C mismatch was positioned 20 nt from the 3’-end of the gRNA^15^. For DECOR, the gRNA was designed to anneal 15 nt from the target edit site based on precedent^15^. For all plasmid cloning, crude ligated product was transformed into DH5α *E. coli*, liquid cultures were grown from single colonies, and miniprepped (Qiagen). Purified plasmids were validated by Sanger sequencing (Genewiz – Azenta Life Sciences).

### Western blotting

To confirm protein expression and analyze formation of editor-ASO covalent complexes in cells, cells were induced with tetracycline (1 µg/mL) for 24 h followed by ASO transfection or no treatment. Cells were harvested and lysed in NP-40 lysis buffer (50 mM Tris HCl pH 7.5, 150 mM NaCl, 5LmM MgCl_2_, 0.5% NP-40, 1 mM PMSF, Roche protease inhibitor tablet). Proteins were separated by SDS-PAGE and analyzed by western blot with anti-HA antibody (ProteinTech, #51064-2-AP, 0.65 µg/mL, 1:5,000-1:2,500 in BSA).

### Synthesis of chloroalkane-modified antisense oligonucleotides (Halo-ASOs)

NH_2_-modified ASOs were synthesized on an ABI 394 oligonucleotide synthesizer (Applied Biosystems) using standard RNA synthesis conditions except for the phosphoramidite coupling time, which was reduced from 12 min to 6 min as recommended for 2’-MOE phosphoramidites. All oligo synthesis reagents were purchased from Glen Research. ASOs were synthesized on Glen Unysupport 500 (Glen Research 20-5140-41) CPG resin unless otherwise indicated (Supplementary Information, Supplementary Table 1), and a primary amine handle was installed on the 5’ end of the ASOs with the 5’-amino-5 modifier (Glen Research 10-1905-90) or internally with the amino modifier C2-dT (Glen Research 10-1037-90). Prior to coupling, the final product was detritylated manually by flowing 3% dichloroacetic acid through the column for 2 min. Oligos were then rinsed with ACN and dried under N_2_. Oligos were cleaved from the CPG resin and deprotected with a 1:1 mixture of NH_4_OH and aq. 40% MeNH_2_ (AMA) for 1 h at 65 °C, following the manufacturer’s recommendations. The supernatant was collected and dried by speedvac. The material was redissolved in 0.1 M triethylammonium acetate (TEAA) and purified via reverse-phase HPLC (0-30% acetonitrile/0.1 M TEAA gradient). The product was lyophilized and redissolved in DEPC-H_2_O. 5’-NH_2_-modified ASOs were coupled to NHS-Halo ligand (Supplementary Information) as follows: to approximately 200 nmol of ASO was added buffer containing 0.5 M MOPS pH 8, 0.5 M NaCl and 0.1 M NHS-Halo ligand in DMSO to give a final concentration of 0.2 M MOPS pH 8, 0.2 M NaCl, and 10 mM NHS-halo ligand. Reactions were carried out for 2 hr at room temperature, diluted with 0.1 M TEAA, purified by RP-HPLC (0-30% acetonitrile in 0.1 M TEAA), and lyophilized. Purified product was dissolved in DEPC-H_2_O, analyzed by ESI-TOF MS (Agilent 6545XT), and stored at -20C.

### TadA8.20-HaloTag-HA expression and purification

pET28a-6xHis-TadA8.20-HaloTag-HA was transformed into *E. coli* BL21 and expressed at 16 °C with 0.1 mM isopropyl-B-D-thiogalactopyranoside (IPTG) for 20 hr. Lysis by sonication was performed in buffer containing 50 mM Tris pH 7.5, 300 mM NaCl, 10 mM imidazole, 10% glycerol, 1 mM DTT, 1 mM PMSF (added fresh), and protease inhibitor tablet (Roche, added fresh). Lysate was clarified at 14,000 *g* for 30 min. Clarified lysate was incubated with 2 mL of fresh HisPur cobalt resin (Thermo Fisher) for 1 hr at 4 °C, washed with lysis buffer, and eluted in the same buffer but with 250 mM imidazole. Fractions were measured by Bradford assay and analyzed by SDS-PAGE, pooled, and dialyzed overnight against 2 L of 50 mM Tris pH 7.5, 100 mM NaCl, 10% glycerol, and 1 mM DTT. Dialyzed protein was then purified via FPLC (GE Healthcare) using a HiTrap QHP 1 mL anion exchange column (100 mM to 1 M NaCl in 50 mM Tris pH 7.5, 10% glycerol, and 1 mM DTT) over 30 CV at 0.5 mL/min. Fractions were analyzed by SDS-PAGE, pooled, and concentrated in 50 mM Tris pH 7.5, 100 mM NaCl, 10% glycerol, 1 mM DTT to > 1 mg/mL, aliquoted at 58.5 μM as determined by Bradford assay, flash frozen, and stored at -80 °C.

### RNA editing with Halo-ASOs in deaminase-HaloTag-expressing Flp-In 293 cells

Culture plates were first coated with poly-L-lysine by diluting 10X poly-L-lysine (Sigma) into 1X DPBS (Gibco Life Technologies) and incubating culture wells for 5 min at RT, aspirating, and washing the wells with 1X DPBS. Wells were allowed to dry for 30 min before plating cells. Editor-HaloTag-expressing Flp-In 293 cells were seeded in a 24-well plate at 7 x 10^4^ cells in 500 μL media. After 24 h, 1 μg/mL tetracycline was added to induce protein expression. After 24 hours, the media was changed, and fresh tetracycline was supplied. Cells were transfected with Halo-ASO (25 nM final concentration in media) together with pcDNA5 plasmid encoding the mRNA reporter (200 ng) unless otherwise indicated. Transfection mixtures were prepared by diluting reagents in OptiMEM with 1.25 µL Lipofectamine 2000 diluted in a separate tube. The solutions were mixed and incubated for 20 min before adding to the wells. Cells were harvested 24 h after transfection unless otherwise indicated.

### RNA editing by lipofection of the TadA8.20-HaloTag-ASO complex

HEK293T or HeLa cells were first plated in a poly-L-lysine-coated 24-well plate at 7.5 x 10^4^ cells/well in 500 µL 24 h before transfection. Shortly before transfection, purified TadA8.20-HaloTag-HA protein was incubated with an equimolar (10 μM) concentration of Halo-ASO in 1X PBS at 37 °C for 30 min. The crude reaction was diluted directly in OptiMEM and 3 μL of Lipofectamine 2000, Lipofectamine 3000, or RNAiMAX was diluted in OptiMEM separately. The solutions were mixed (50 μL total volume) and incubated for 20 min before adding to the wells without changing the media to achieve a final enzyme-ASO complex concentration of 25 nM in 550 μL media. Cells were harvested after 24 h for RNA analysis.

### RNA editing by electroporation of the TadA8.20-HaloTag-ASO complex

HEK293T cells were grown to 80% confluency before being trypsinized, pelleted, and resuspended in OptiMEM to a concentration of 5.556 x 10^6^ cells/mL. TadA8.20-HaloTag-ASO complex was prepared freshly as described above, and 10 μL of the reaction was mixed with 90 μL of cell suspension to achieve a 1 μM enzyme-ASO complex concentration in 100 μL with 5 x 10^5^ cells. The mixture was transferred to a 2 mm electroporation cuvette (Universal Medical) and electroporated (100 – 160 V, 10 ms or 20 ms time constant) using a BTX ECM 830 device. Cells were allowed to recover for 10 min at room temperature before mixing with 1.9 mL culture media and plating in a 6-well plate. Cells were harvested after 24 h for RNA analysis.

### RNA isolation and Sanger sequencing analysis

RNA isolation was performed with TRIzol (Thermo Fisher), TRI Reagent (Molecular Research Center), or RNAzol RT (Molecular Research Center) following the manufacturer’s protocol but with 250-500 μL reagent per sample and proportionally scaled volumes of the other reagents utilized in the procedure. After isopropanol precipitation and washing with 75% ethanol, the RNA pellets were dissolved in DEPC-treated water, DNase buffer, and treated with DNase I (NEB) or TURBO DNase (Thermo Fisher) for 30 min at 37 °C before ethanol precipitation at -80 °C. Samples were then pelleted by centrifugation, washed with 70% ethanol, and resuspended in DEPC water. The concentration was then determined by Nanodrop (Thermo Fisher). 1-2 μg of total RNA was used in the reverse transcription reactions using SuperScript II (Invitrogen) or in-house-purified MMLV^81^ (Addgene #153312) following the Invitrogen SuperScript II protocol. Generally, 4 μL of crude reverse transcription reaction was used as the template in a 25 μL PCR reaction employing Taq DNA polymerase (NEB). Products were analyzed by agarose gel electrophoresis and gel purified before Sanger sequencing (Genewiz – Azenta Life Sciences). EditR^51^ (https://moriaritylab.shinyapps.io/editr_v10/) was used to quantify editing stoichiometry from Sanger sequencing data, and a student’s t-test was used to assess statistical significance.

### RNA editing with REPAIRv2 and DECOR

HEK293T cells were seeded on a poly-L-lysine-coated 12-well plate at 1.3 x 10^5^ cells/well 24 h before transfection. For REPAIRv2, 750 ng of editor expression plasmid (Addgene #103871) and 375 ng of gRNA-expressing plasmid was diluted to 125 μL in OptiMEM and mixed with 2.5 μL Lipofectamine 2000 diluted to 125 μL separately in OptiMEM. Complexes were incubated for 20 min and added to fresh 750 μL media. In the case of PTEN Q245X-directed editing, 200 ng of PTEN Q245X plasmid was also included in the transfection. The conditions were the same for DECOR, except that 500 ng DECOR expression plasmid (Addgene #219545) and 750 ng gRNA-expressing plasmid were used. Cells were harvested after 24 h for RNA isolation and sequencing analysis as described above.

### RNA-seq library preparation

HEK293T cells were plated on poly-L-lysine-coated 6-well plate at 2.75 x 10^5^ cells/well 24 h before transfection. RNA-protein complexes were formed *in vitro* as described above. The crude reaction was diluted directly in OptiMEM, and 12 μL of Lipofectamine 2000 was diluted in OptiMEM separately. The solutions were mixed (200 μL total volume) and incubated for 20 min before adding to the wells without changing the media to achieve a final enzyme-ASO complex concentration of 25 nM in 2.2 mL media. RNA was isolated after 24 h using TRI Reagent (Molecular Research Center) and treated with DNase I (New England Biolabs) prior to mRNA isolation via polyA pulldown with oligo dT(25) magnetic beads (New England Biolabs). 50 ng of polyA mRNA was subject to library preparation using the NEBNext Ultra II Directional RNA Library Prep Kit for Illumina (New England Biolabs). Quality control was determined by gel electrophoresis and Bioanalyzer. The libraries were sequenced on an Illumina NovaSeq 6000 instrument (paired-end; R1 80 nt, R2 250 nt).

### Bioinformatic analysis

Sequences were demultiplexed based on the barcoded-index primers using barcode-splitter. Demultiplexed raw reads were then uploaded to the Princeton Della HPC cluster for further analysis. The sequencing data for other RNA-editing methods were obtained from Gene Expression Omnibus (GEO) using SRA Toolkit. The analysis pipeline and scripts were obtained and adapted from Yan and Tang^15^. Briefly, the R3 reads were trimmed to 97 bp using Cutadapt and aligned to the hg19 (GRCh37) reference genome using STAR 2.7.11b. Unmapped reads were discarded and PCR duplicates were removed using SAMtools 1.21. Deduplicated BAM files were randomly down-sampled to 7.4 million reads to ensure uniform sequencing depth across samples using BBTools. A-to-G mutations were called using REDItools2.0. Only A sites with ≥10 reads in the control and ≥10 reads in the test condition were retained. A SNP table was not used for filtering SNPs. Instead, editing values >90% in the control samples were considered SNPs and were filtered out. A Fisher’s exact test was performed in test and control cells to determine statistically significant A-to-G or T-to-C (depending on whether forward or reverse reads were used) edit sites. Sites with P > 0.05 were removed. The overlap of edit sites was visualized with MetaChart. Sequence motif analysis was performed with MEME in The MEME Suite 5.5.8.

## SUPPORTING INFORMATION

Purification of recombinant proteins, small molecule synthesis, oligonucleotide characterization, protein-oligo crosslinking reactions, supporting data for RNA editing reactions, and whole-transcriptome RNA editing analysis.

## ACKNOWLEDGEMENTS

R.E.K. acknowledges support from the NIH (R01 GM152748) and NSF (MCB-1942565). T.W.E. and D.Y.D. were supported by a generous gift from the Edward C. Taylor 3rd Year Graduate Fellowship in Chemistry. All authors acknowledge financial support from Princeton University.

