## Supplementary Information for "Targeted RNA editing by direct delivery of an adenosine deaminase-antisense oligo conjugate"

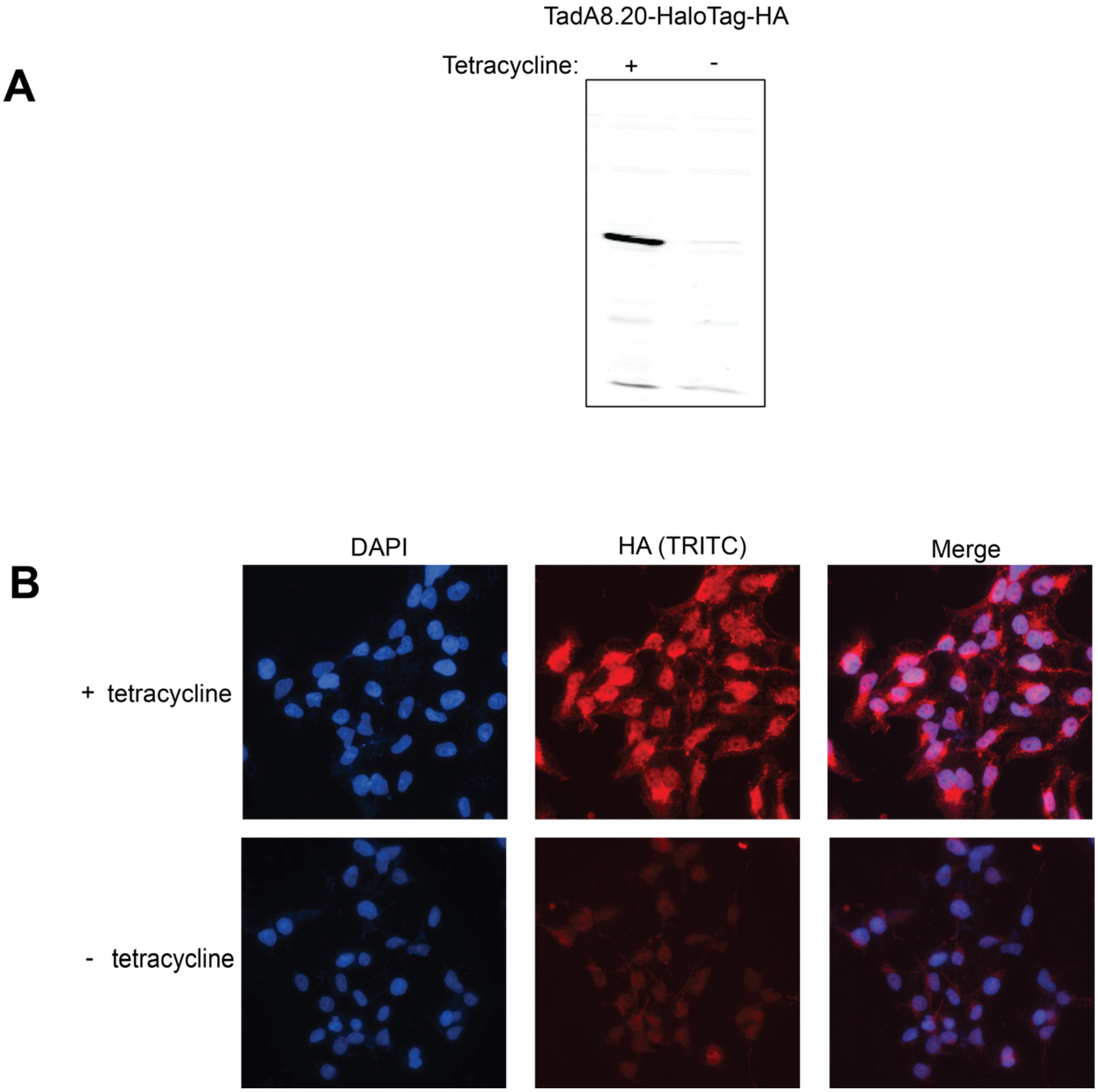


**Supplementary Figure 1.** Analysis of protein expression in TadA8.20-HaloTag Flp-In TREx 293 cell line. **(A)** Western blot showing expression of TadA8.20-HaloTag expression after 24 h induction with 1 µg/mL tetracycline. **(B)** Immunofluorescence microscopy of TadA8.20-HaloTag-HA.

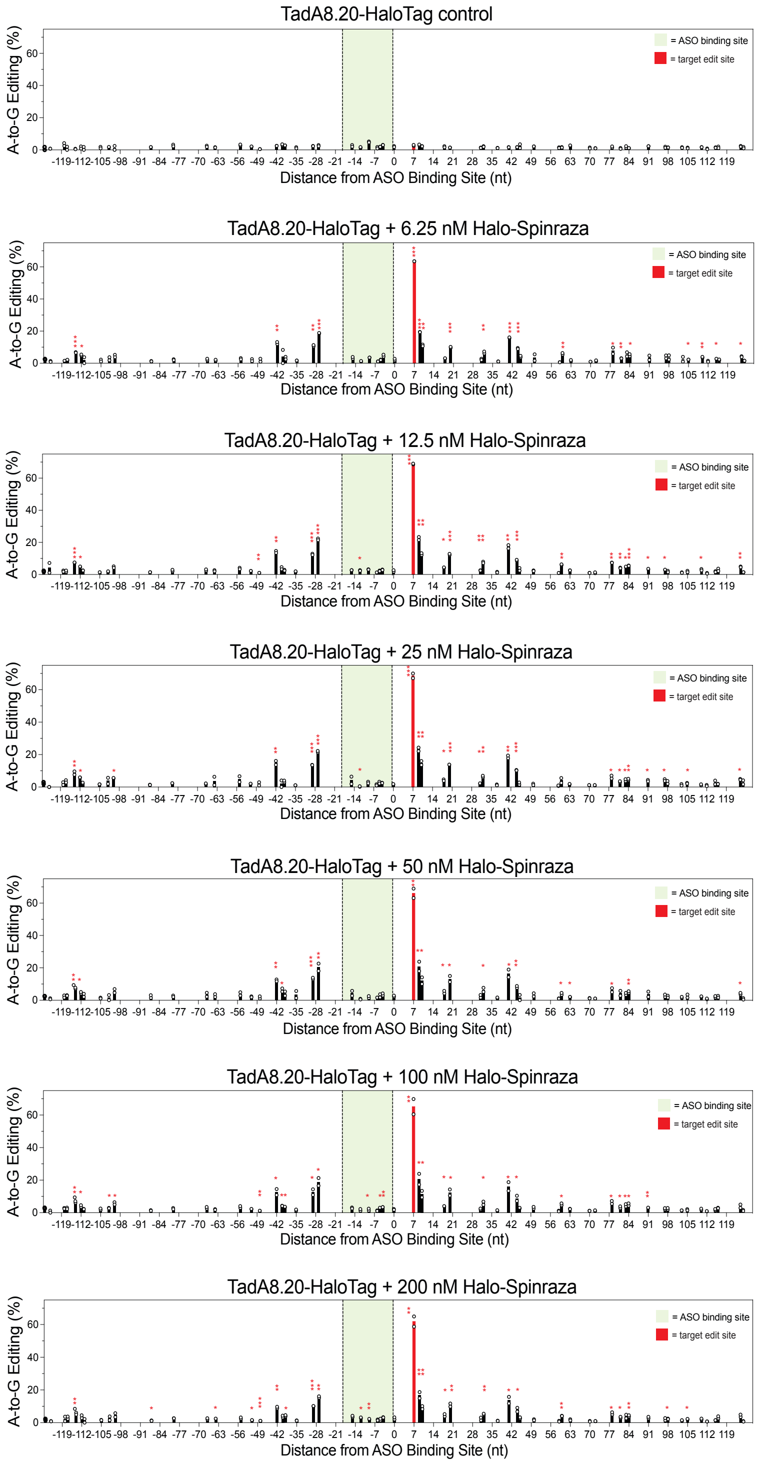


**Supplementary Figure 2.** TadA8.20-HaloTag + Halo-Spinraza 1 dose titration editing. The data shown in the heatmap in Figure 1d is shown here for each tested concentration. Experiments were performed in TadA8.20-HaloTag Flp-In 293 cells. At least two independent biological replicates were performed for all experiments. Statistical significance legend: * = p ≤ 0.05, ** = p ≤ 0.01, *** = p ≤ 0.001.

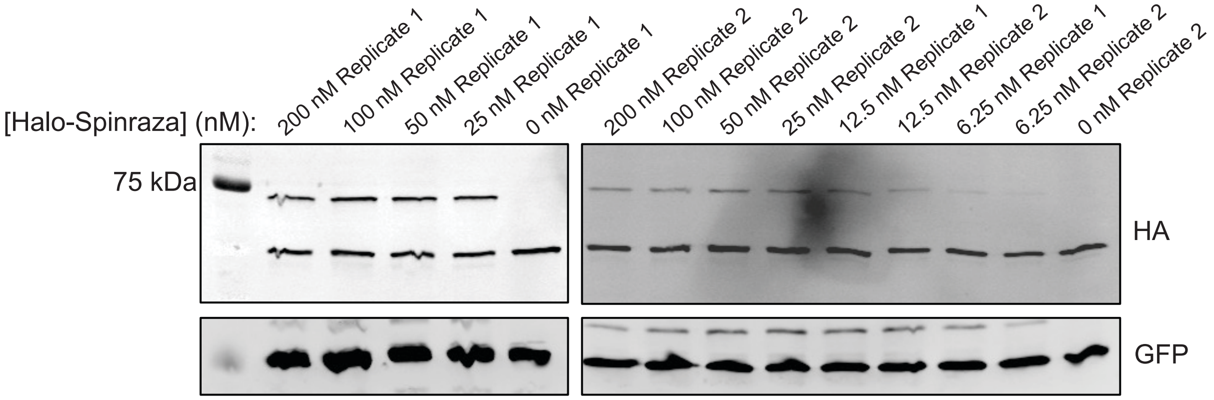


**Supplementary Figure 3.** Halo-Spinraza 1 crosslinking at each tested ASO concentration. Halo-Spinraza 1 was transfected into TadA8.20-HaloTag Flp-In 293 cells and harvested after 24 h. Two independent biological replicates were performed for all experiments.

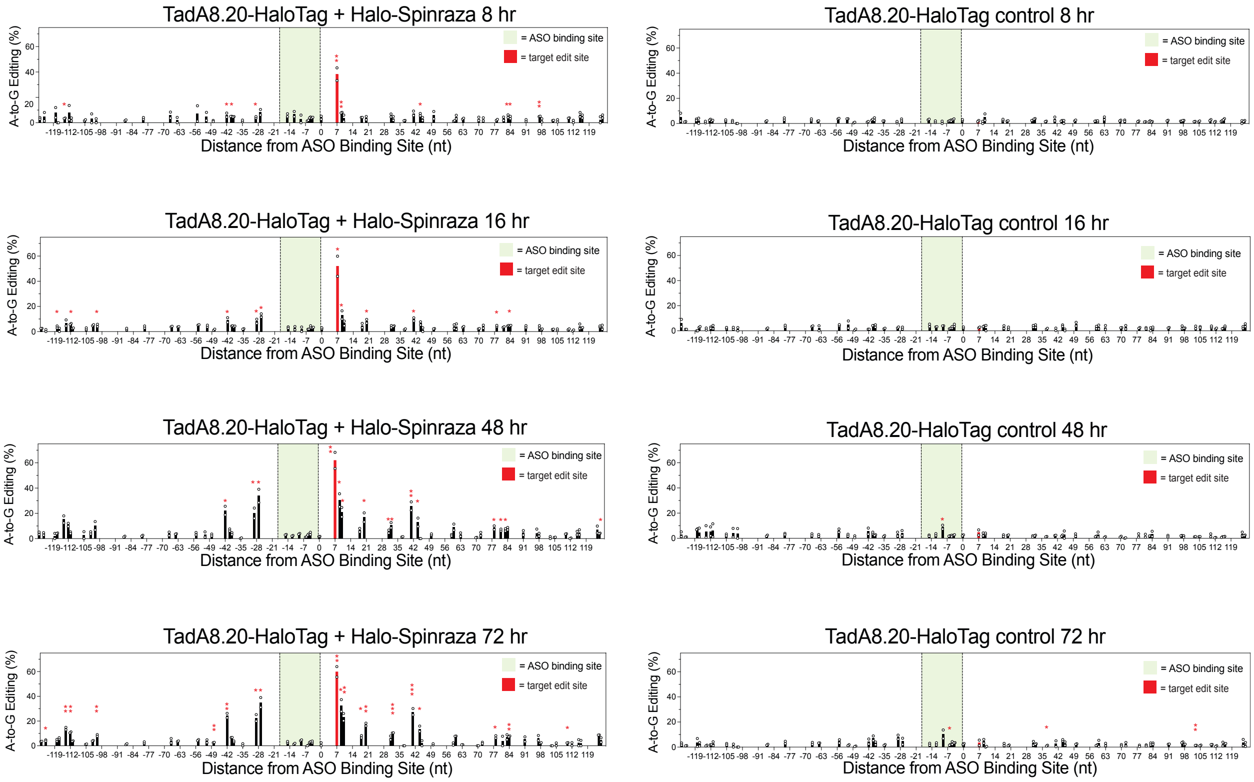


**Supplementary Figure 4.** TadA8.20-HaloTag + Halo-Spinraza 1 time course editing. The data shown in the heat map in Figure 1e is shown here as individual plots for each time point along with the no ASO control for each. Experiments were performed in TadA8.20-HaloTag Flp-In 293 cells. Two independent biological replicates were performed for all experiments. Statistical significance legend: * = p ≤ 0.05, ** = p ≤ 0.01, *** = p ≤ 0.001.

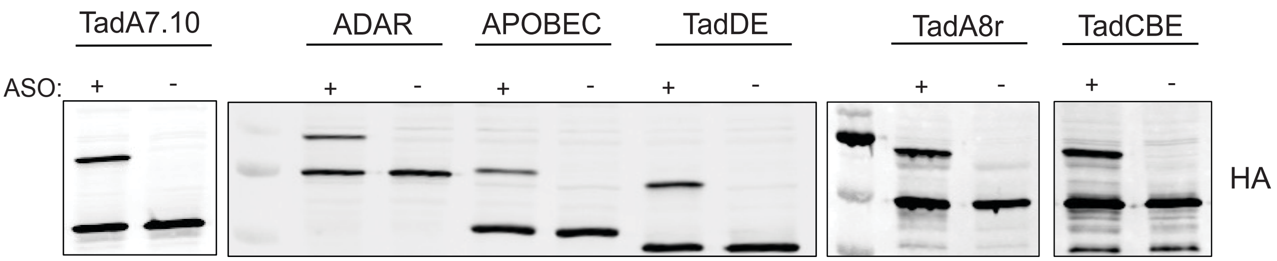


**Supplementary Figure 5.** Western blot analysis of in-cell conjugation between Halo-Spinraza 1 and RNA editor-HaloTag fusion proteins in RNA editor-HaloTag Flp-In 293 cells.

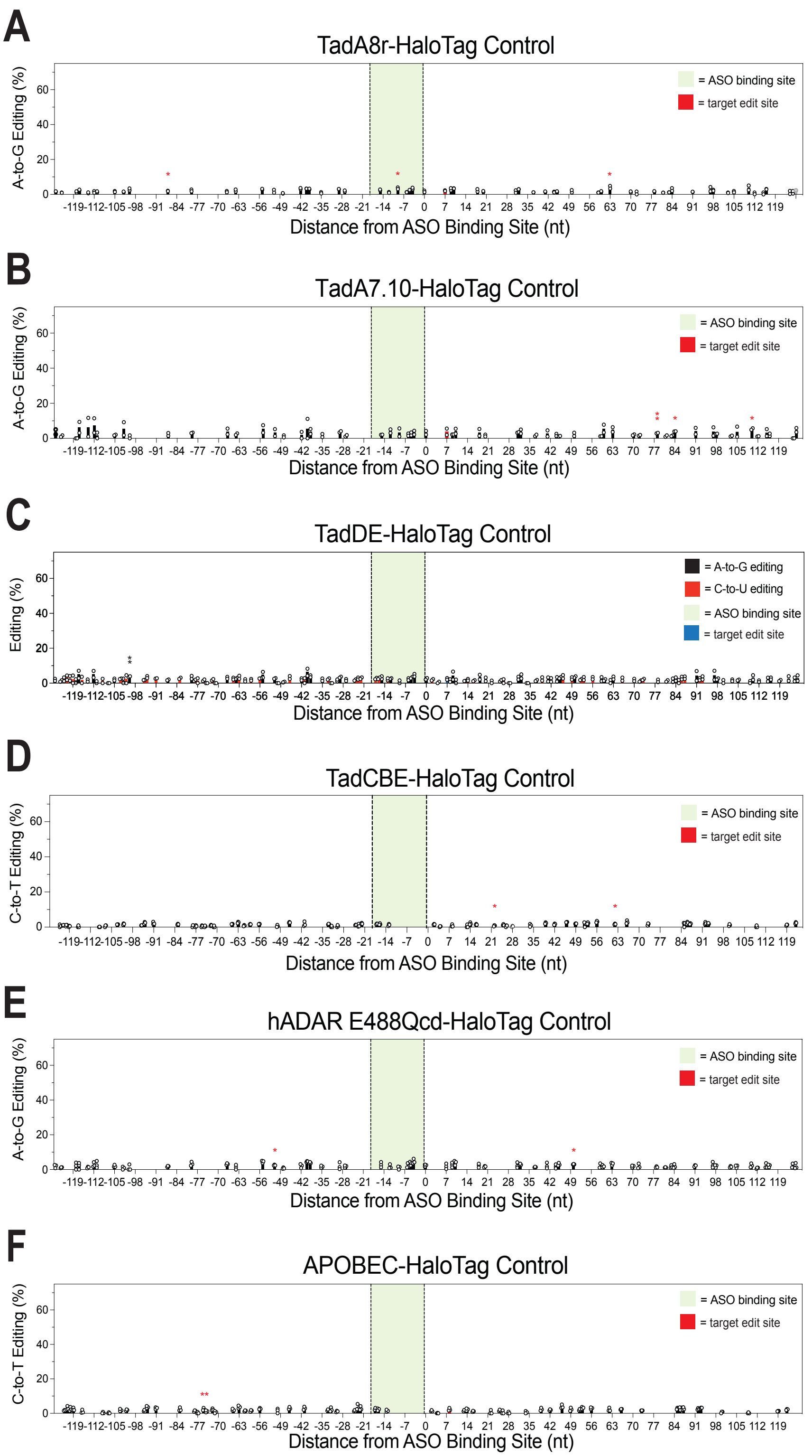


**Supplementary Figure 6.** Negative control (no ASO) editing data for HaloTag-fusion proteins. Experiments were performed in TadA8.20-HaloTag Flp-In 293 cells. At least two independent biological replicates were performed for all experiments. Statistical significance legend: * = $p \leq 0.05$, ** = $p \leq 0.01$, *** = $p \leq 0.001$.

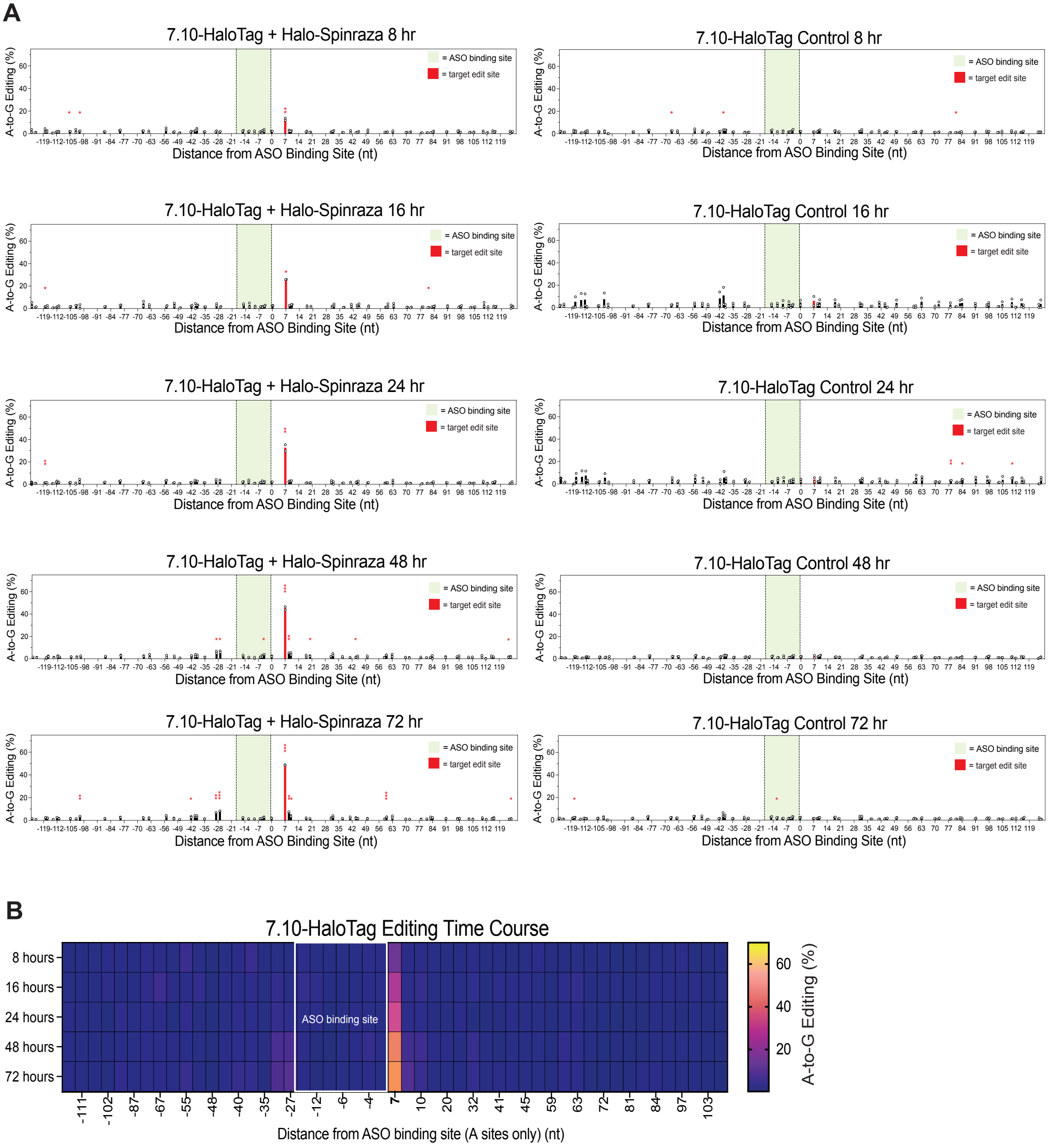


**Supplementary Figure 7. (A)** TadA7.10-HaloTag + Halo-Spinraza time course editing. **(B)** Heat map representing the data shown in **(A)**. Experiments were performed in TadA8.20-HaloTag Flp-In 293 cells. At least two independent biological replicates were performed for all experiments. Statistical significance legend: * = p ≤ 0.05, ** = p ≤ 0.01, *** = p ≤ 0.001.

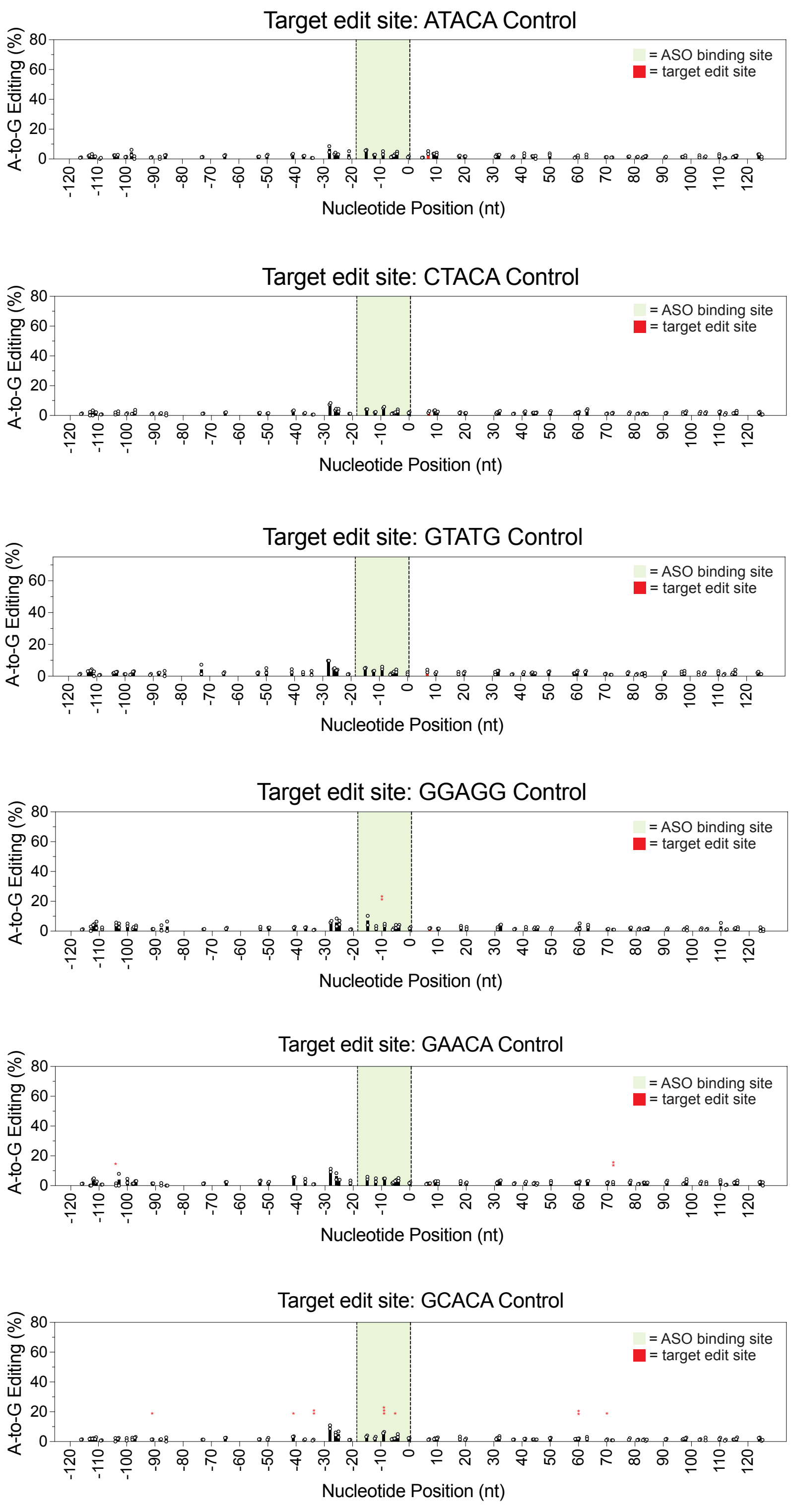


**Supplementary Figure 8.** Negative control editing data for sequence motif mutants in Figure 3. Experiments were performed in TadA8.20-HaloTag Flp-In 293 cells. Two independent biological replicates were performed for all experiments. Statistical significance legend: * = p ≤ 0.05, ** = p ≤ 0.01, *** = p ≤ 0.001.

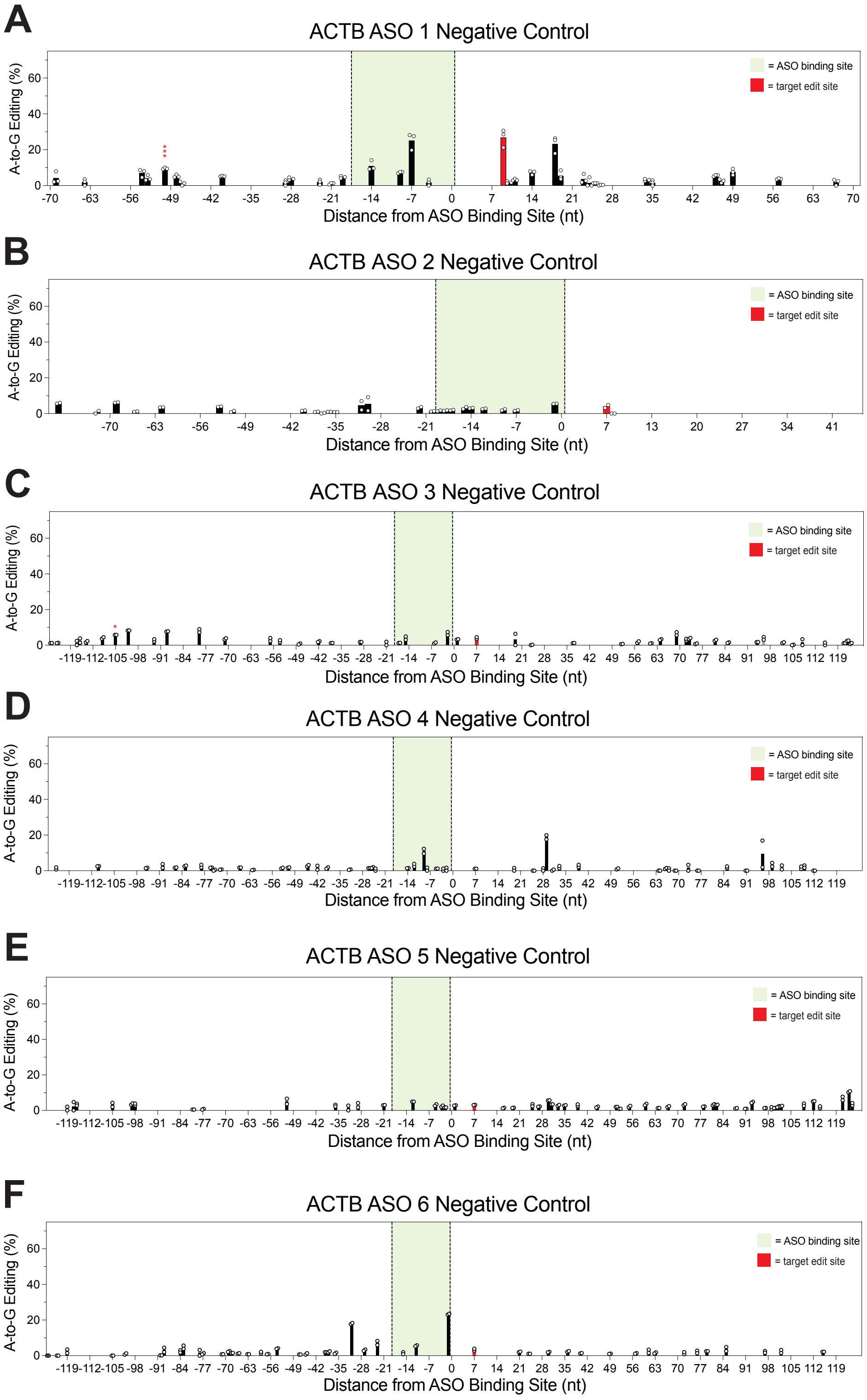


**Supplementary Figure 9.** Negative controls (no ASO) for ACTB ASOs 1-6 in Figure 4. Experiments were performed in TadA8.20-HaloTag Flp-In 293 cells. At least two independent biological replicates were performed for all experiments. Statistical significance legend: * = $p \leq 0.05$, ** = $p \leq 0.01$, *** = $p \leq 0.001$.

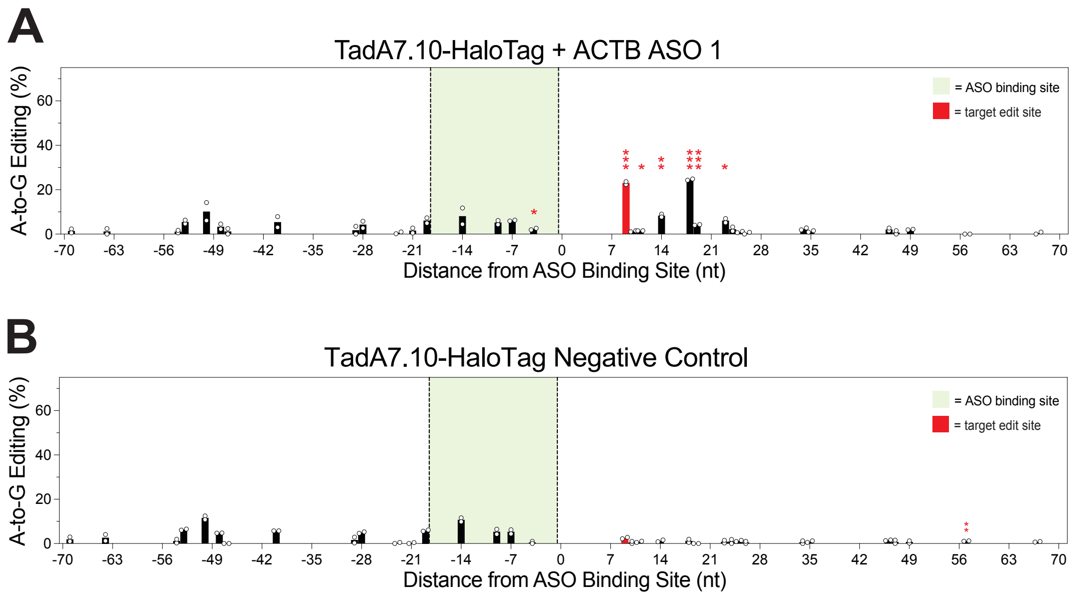


**Supplementary Figure 10.** Editing on endogenous ACTB 3'-UTR using ACTB ASO 1 transfected into cells expressing TadA7.10-HaloTag. Two independent biological replicates were performed for all experiments. Statistical significance legend: * = $p \leq 0.05$, ** = $p \leq 0.01$, *** = $p \leq 0.001$.

**A**

```
ACTB   601 CCACCACGGCCGAGCGGAAATCGTGCGTGACATTAAGGAGAAGCTGTGC    650
           |||||||||||||||||||||||||||.|||||.||||||||||||||||
ACTG1  574 CCACCACGGCCGAGCGGAAATCGTGCGCGACATCAAGGAGAAGCTGTGC    623

ACTB   651 TACGTCGCCCTGGACTTCGAGCAAGAGATGGCCACGGCTGCTTCCAGCTC    700
           |||||||||||||||||||||||.|||||||||||.||.||.|||..|||
ACTG1  624 TACGTCGCCCTGGACTTCGAGCAGGAGATGGCCACCGCCGCATCCTCCTC    673
```

**B**

```
ACTB   450 catgtttgagaccttcaacaccccagccatgtacgttgctatccaggctg    499
           .|||||||||||||||||||||||.|||||||||||.||.||||||||.|
ACTG1  375 tatgtttgagaccttcaacaccccggccatgtacgtggccatccaggccg    424

ACTB   500 tgctatccctgtacgcctctggccgtaccactggcatcgtgatggactcc    549
           ||||.|||||.|||||||||||.||.|||||||||||.||.||||||||.
ACTG1  425 tgctgtccctctacgcctctgggcgcaccactggcattgtcatggactct    474
```

**Supplementary Figure 11.** Sequence alignment for ACTB and ACTG1. **(A)** Alignment for the region around the ACTB ASO 4 binding site. **(B)** Alignment for the region around the ACTB ASO 6 binding site. Alignment of human actin genes was performed with Emboss Needle. A green box denotes the ASO binding site. A-to-G variations between ACTB and ACTG1 that appear in the negative controls in Supplementary Figure 9 are highlighted in yellow.

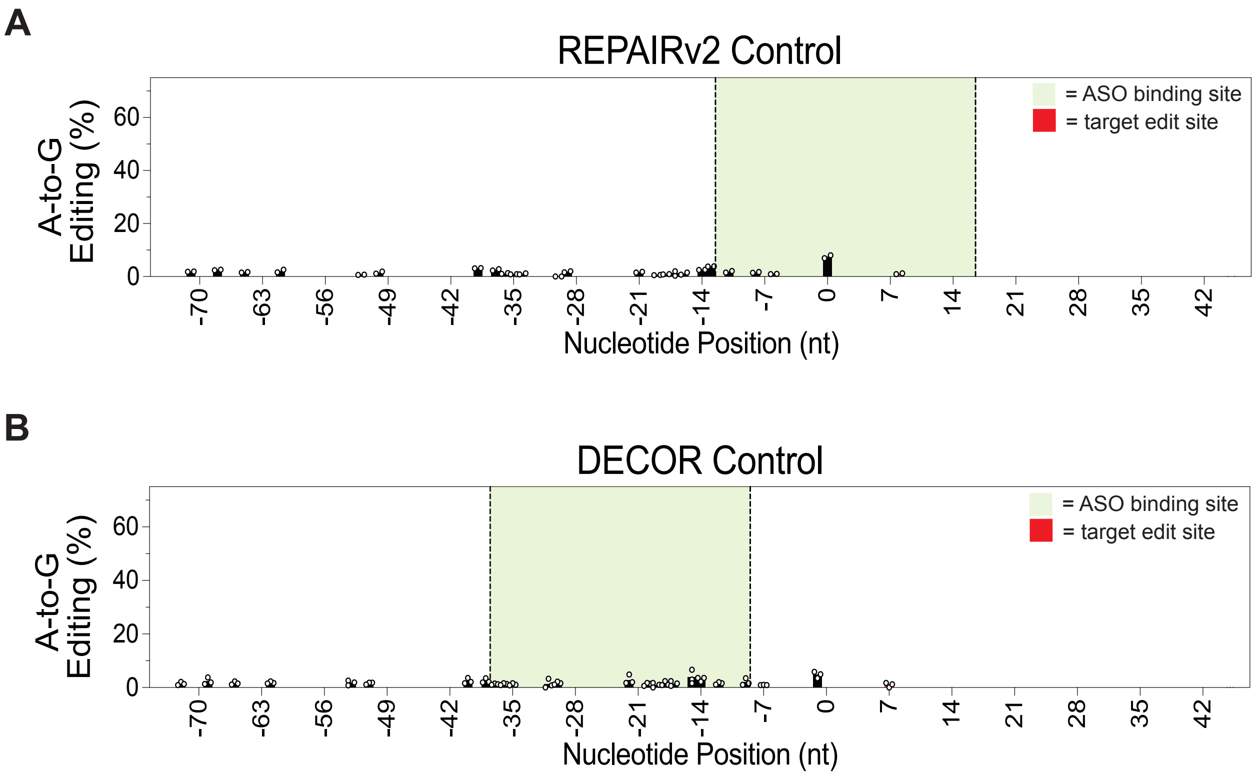


**Supplementary Figure 12.** Negative controls for editing in ACTB 3'-UTR using REPAIRv2 **(A)** and DECOR **(B)**. Two independent biological replicates were performed for all experiments. Statistical significance legend: * = $p \leq 0.05$, ** = $p \leq 0.01$, *** = $p \leq 0.001$.

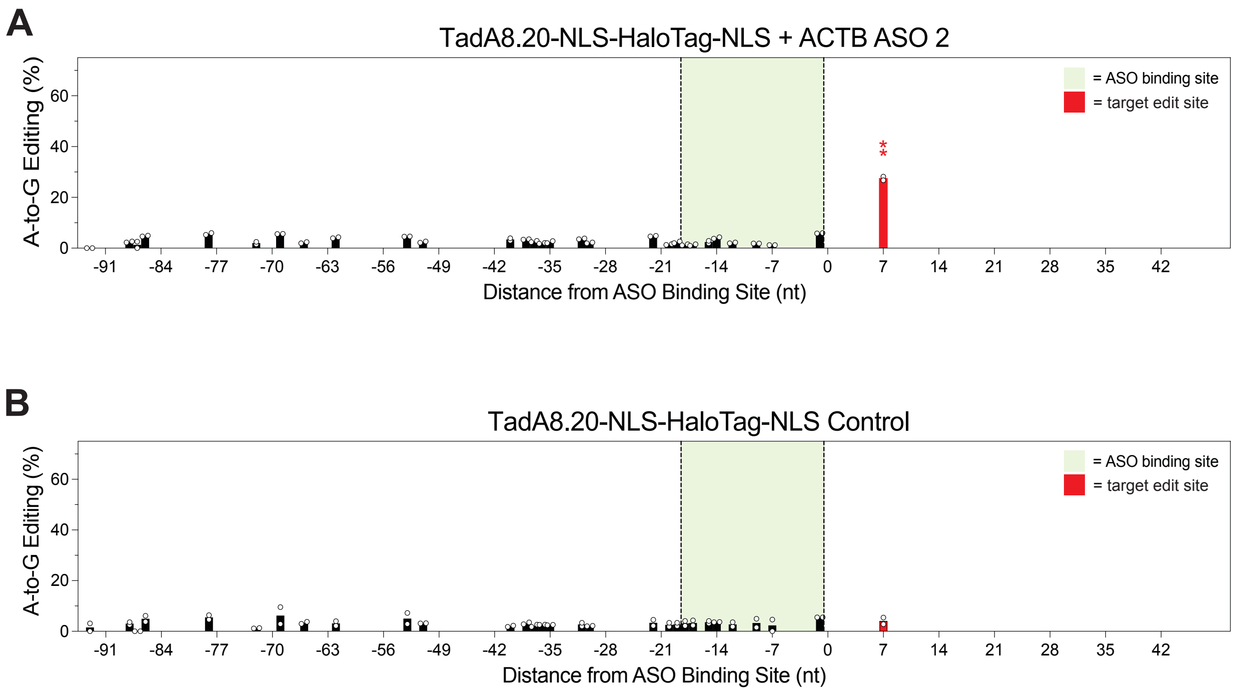


**Supplementary Figure 13.** Editing of ACTB 3'-UTR with ACTB ASO 2 transfected into TadA8.20-NLS-HaloTag-NLS expressing cells. Two independent biological replicates were performed for all experiments. Statistical significance legend: * = $p \leq 0.05$, ** = $p \leq 0.01$, *** = $p \leq 0.001$.

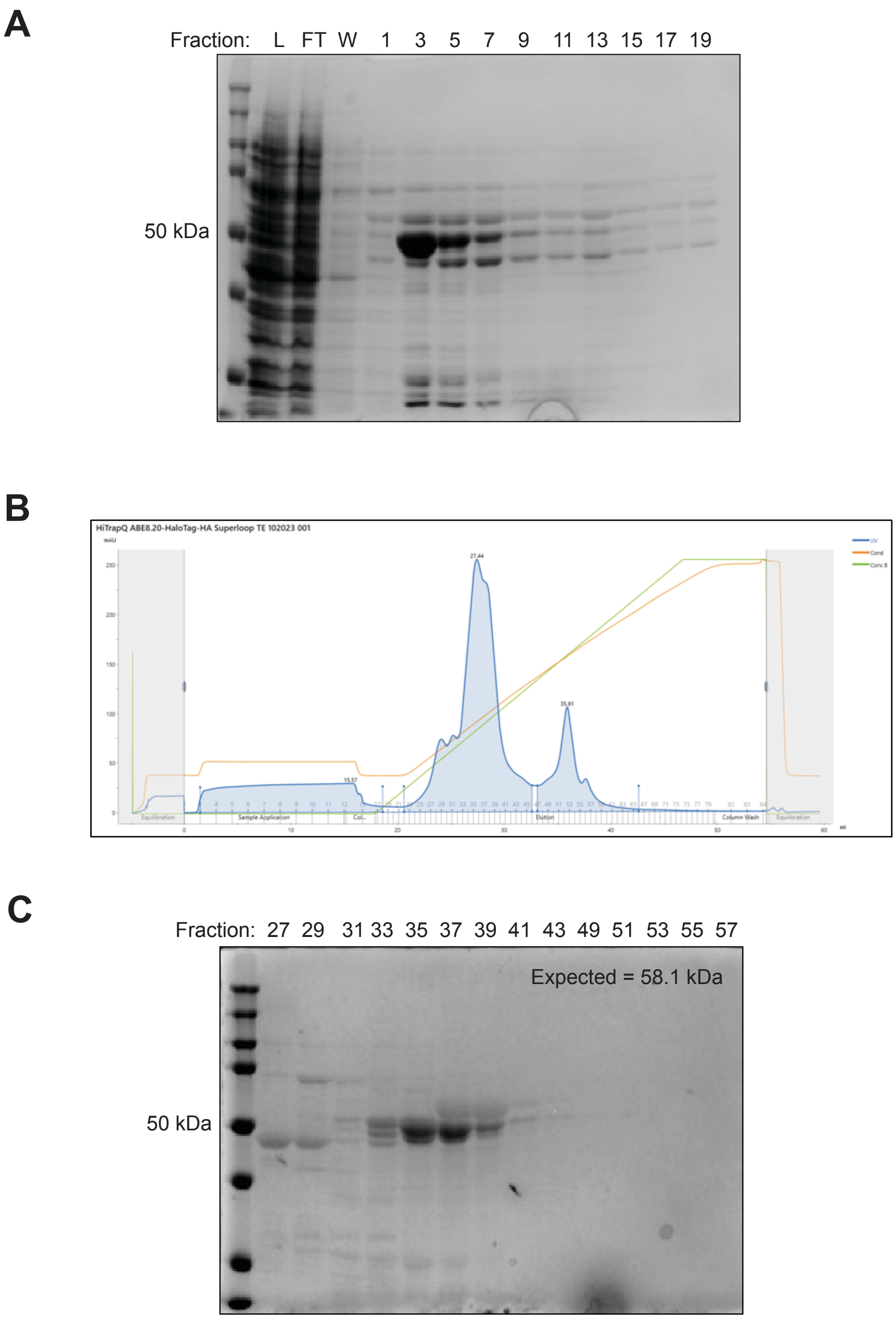


**Supplementary Figure 14.** Purification of recombinant TadA8.20-HaloTag**. (A)** Coomassie-stained SDS-PAGE gel from Co-affinity purification of TadA8.20-HaloTag. Fractions 2-15 were pooled and dialyzed overnight against 2 L of 50 mM Tris pH 7.5, 100 mM NaCl, 10% glycerol, and 1 mM DTT before anion exchange chromatography. **(B)** Anion exchange chromatogram of post-Co affinity purified TadA8.20-HaloTag. **(C)** Coomassie-stained SDS-PAGE gel of fractions 27-57 isolated from the anion exchange chromatography purification shown in **(B)**. Fractions 34-36 were pooled and concentrated.

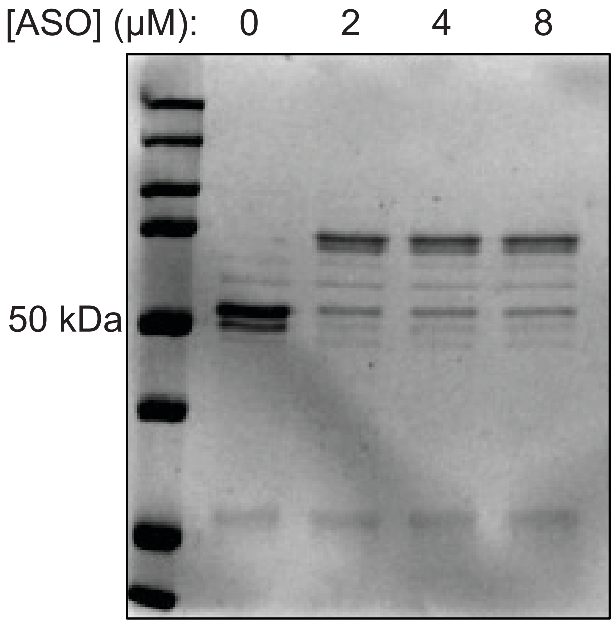


**Supplementary Figure 15.** Purified recombinant TadA8.20-HaloTag-HA conjugation with Halo-Spinraza ASO. Halo-Spinraza ASO was incubated with 2 μM enzyme at a 0:1, 1:1, 2:1, and 4:1 ratio in 1X PBS for 30 min at 37 °C and analyzed by Coomassie-stained SDS-PAGE.

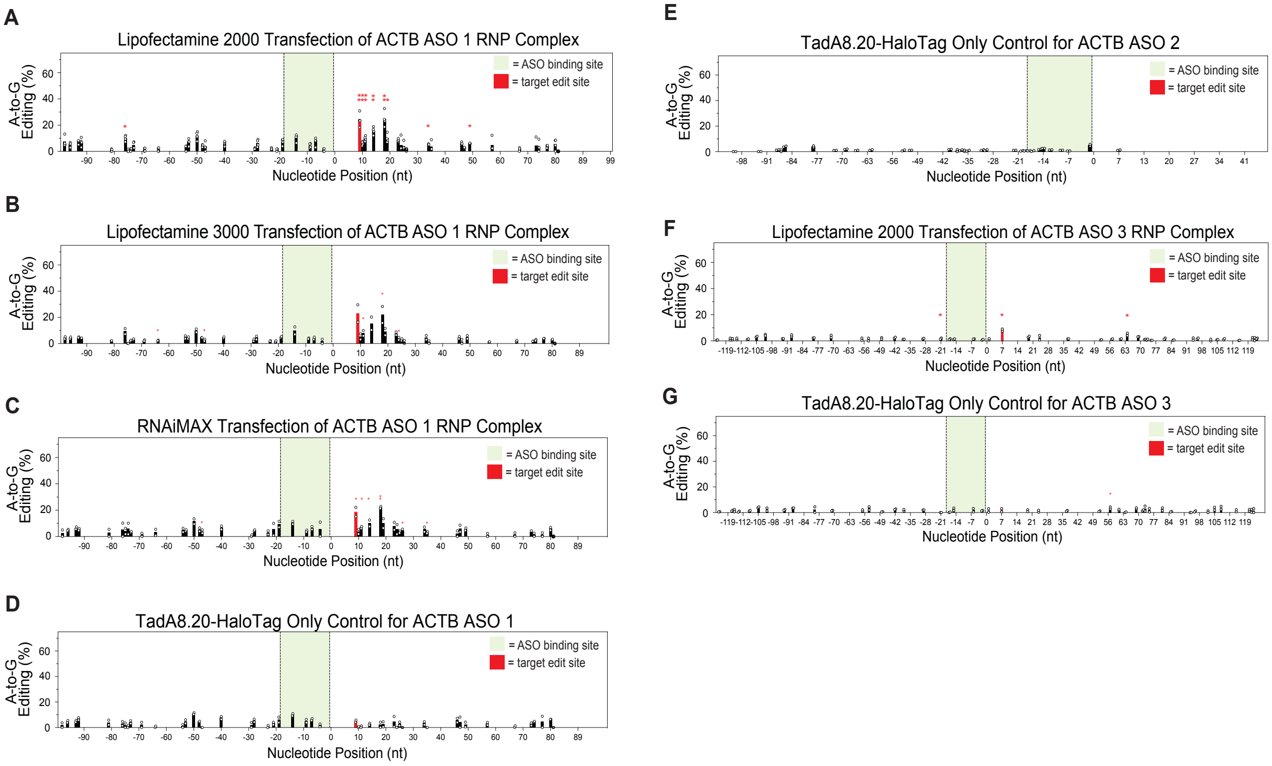


**Supplementary Figure 16.** Lipofection of RNP complexes. Protein-ASO complexes were formed with ACTB ASO 1, 2, or 3 and TadA8.20-HaloTag protein, then transfected into HEK293T cells at 25 nM with Lipofectamine 2000 **(A)**, Lipofectamine 3000 **(B)**, or Lipofectamine RNAiMAX **(C)**. Lipofectamine 2000 was used for **(D-G)**. **(D)** Enzyme only control for ACTB ASO 1. **(E)** Enzyme only control for ACTB ASO 2 in HEK293T cells. **(F)** Transfection of ACTB ASO 3 RNP complex. **(G)** Enzyme only control for ACTB ASO 3 in HEK293T cells. Two independent biological replicates were performed for all experiments. Statistical significance legend: * = $p \leq 0.05$, ** = $p \leq 0.01$, *** = $p \leq 0.001$.

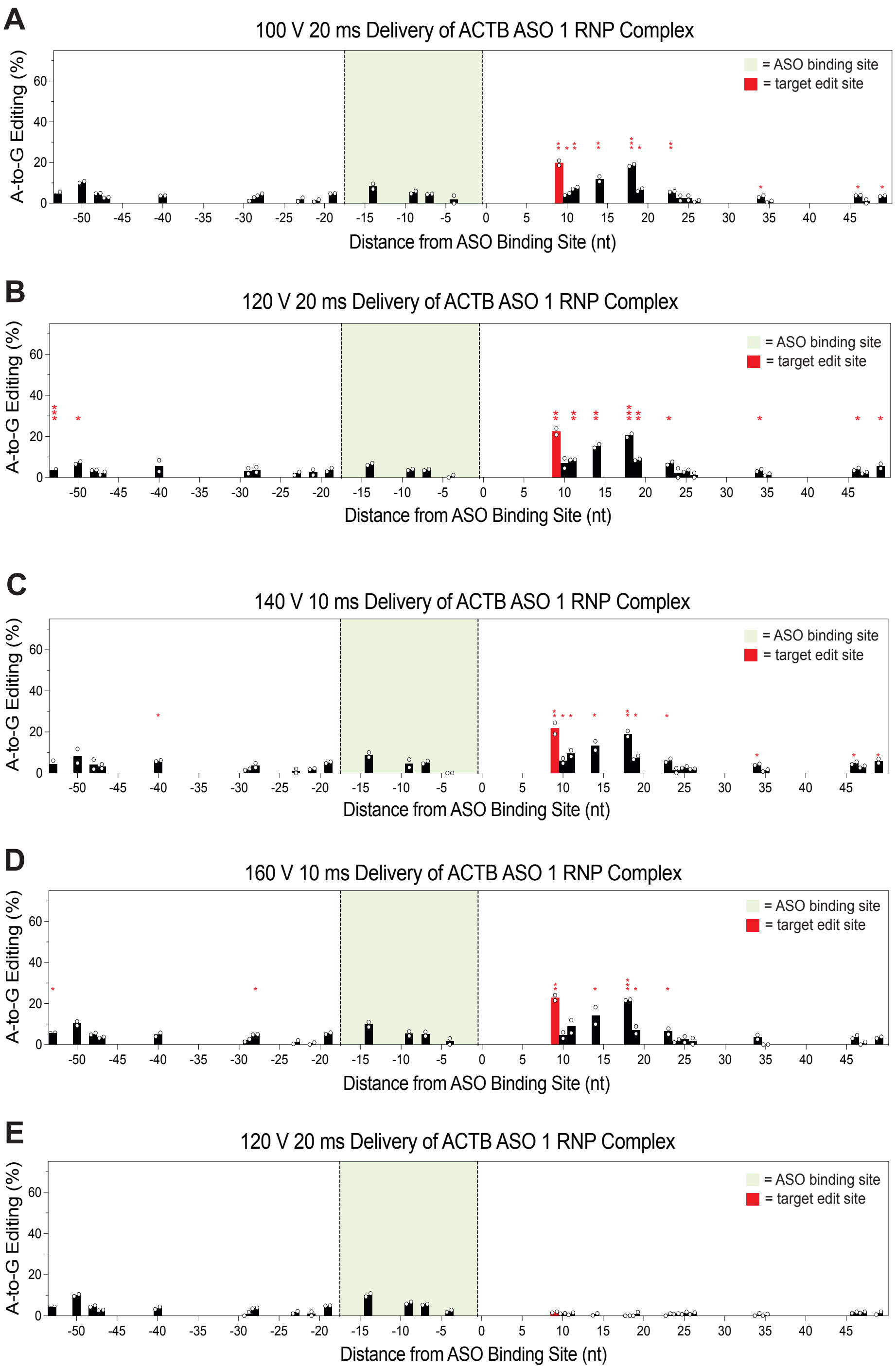


**Supplementary Figure 17.** Electroporation of ACTB ASO 1 RNP complex yields similar editing across tested parameters. **(A)** 100 V, 20 ms electroporation. **(B)** 120 V, 20 ms electroporation. **(C)** 140 V, 10 ms electroporation. **(D)** 160 V, 10 ms electroporation. **(E)** 120 V, 20 ms electroporation of recombinant TadA8.20-HaloTag without conjugated ASO. Cells electroporated at >160 V did not survive (data not shown). Two independent biological replicates were performed for all experiments. Statistical significance legend: * = $p \leq 0.05$, ** = $p \leq 0.01$, *** = $p \leq 0.001$.

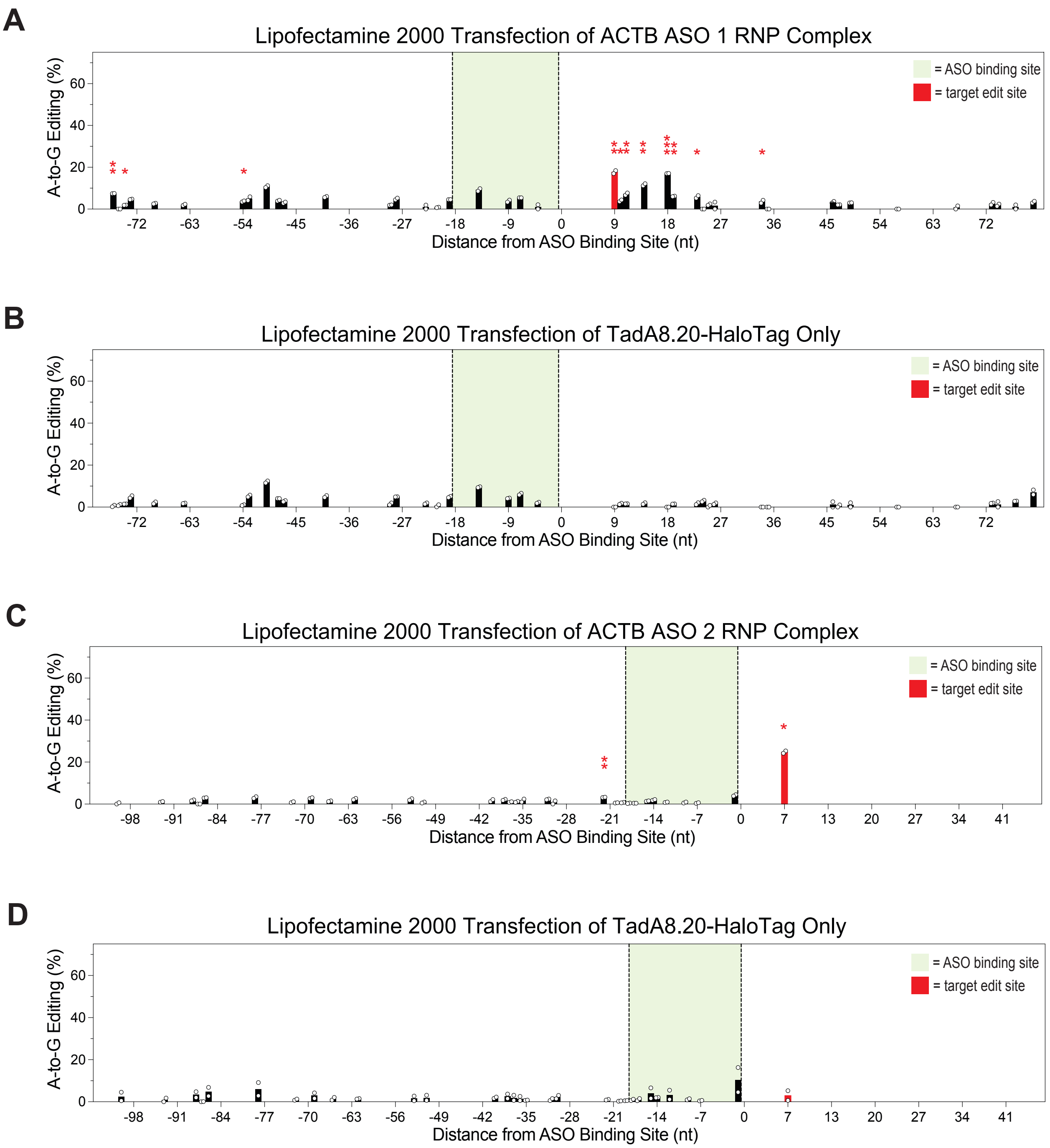


**Supplementary Figure 18.** TadA8.20-HaloTag-ASO complex editing of ACTB mRNA in HeLa cells. **(A)** ACTB ASO 1 RNP complex. **(B)** Enzyme only for ACTB ASO 1. **(C)** ACTB ASO 2 RNP complex. **(D)** Enzyme only for ACTB ASO 2. Two independent biological replicates were performed for all experiments. Statistical significance legend: * = $p \leq 0.05$, ** = $p \leq 0.01$, *** = $p \leq 0.001$.

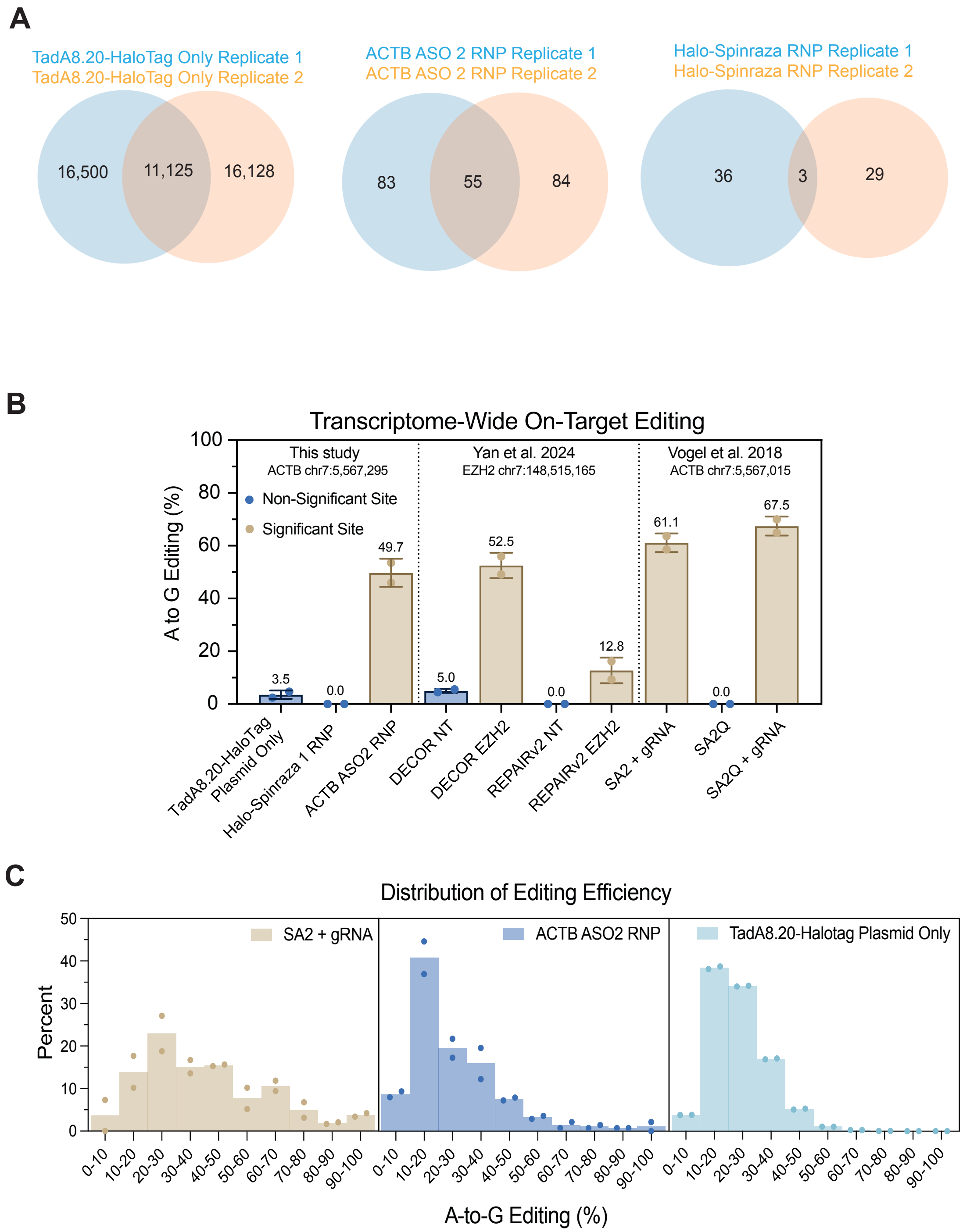


**Supplementary Fig. 19.** Transcriptome-wide editing analysis. **(A)** Overlapping sites between replicates of each editing condition. **(B)** Target site editing percentage for the three tested conditions compared to other published RNA editing techniques. **(C)** Off-target editing rate comparison of SNAP-ADAR2 (SA2) + gRNA (left), ACTB ASO 2 RNP (middle), and TadA8.20-HaloTag plasmid only (right). Histograms show the percentage of detected edit sites in each decile of editing percent range. Bars indicate the mean of two replicates.

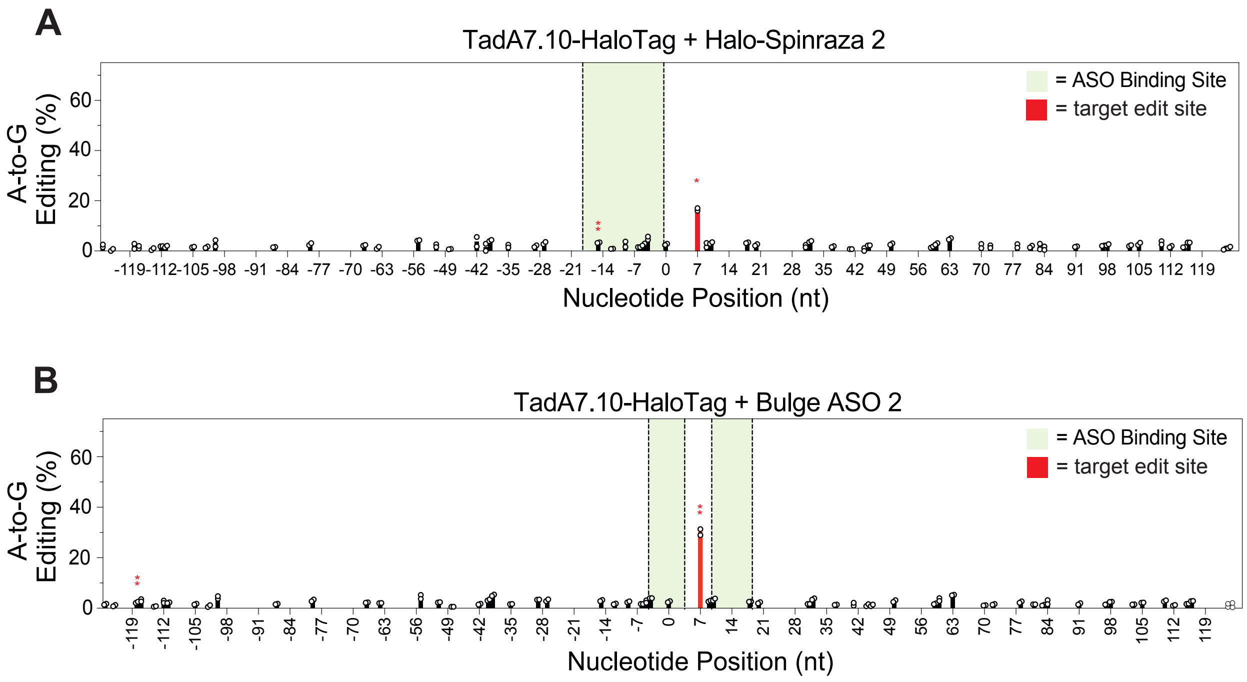


**Supplementary Figure 20.** Editing of dual reporter mRNA in TadA7.10-HaloTag-expressing HEK293 Flp-In cells, transfected with Halo-Spinraza ASO 2 **(A)** or Bulge ASO 2 **(B)**. Statistical significance legend: * = $p \leq 0.05$, ** = $p \leq 0.01$, *** = $p \leq 0.001$.

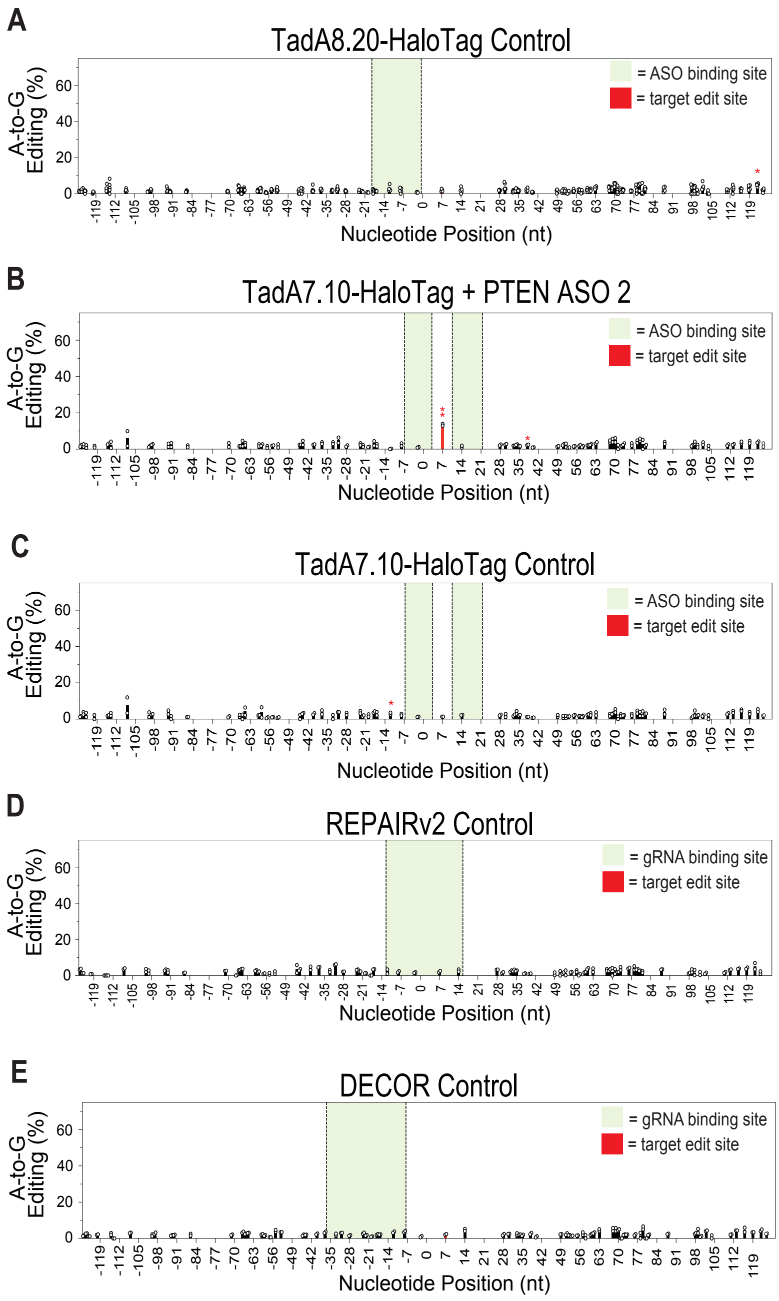


**Supplementary Figure 21.** Editing of the PTEN Q245X mutation. **(A)** No ASO control in TadA8.20-HaloTag-expressing cells. **(B)** Editing using PTEN ASO 2 in TadA7.10-HaloTag-expressing cells. **(C)** No ASO control in TadA7.10-HaloTag-expressing cells. **(D)** REPAIRv2 no gRNA control. **(E)** DECOR no gRNA control. At least two independent biological replicates were performed for all experiments. Statistical significance legend: * = $p \leq 0.05$, ** = $p \leq 0.01$, *** = $p \leq 0.001$.

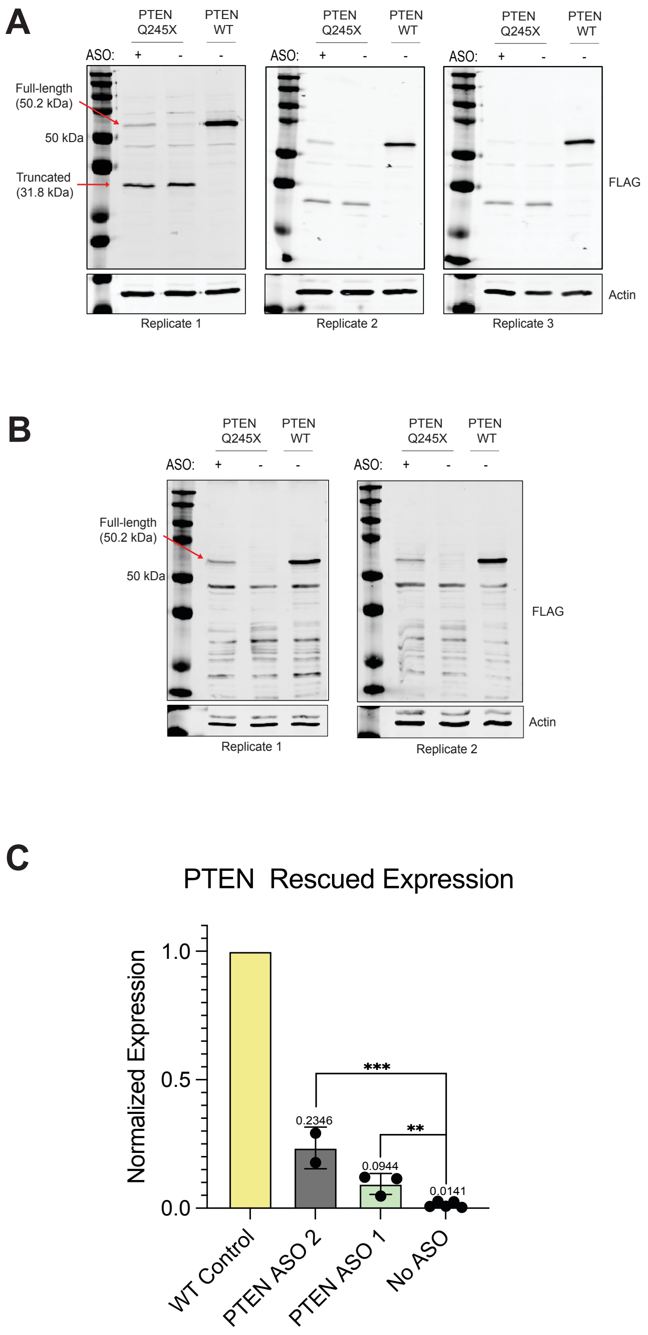


**Supplementary Figure 22.** Quantification of PTEN protein expression after correction of Q245X mutation. **(A)** Western blots 24 h after transfection of PTEN ASO 1 into TadA8.20-HaloTag Flp-In 293 cells. **(B)** Western blots 24 h after transfection of PTEN ASO 2 into TadA8.20-HaloTag Flp-In 293 cells. **(C)** Quantification of restored PTEN expression. At least two independent biological replicates were performed for all experiments. Statistical significance legend: * = $p \leq 0.05$, ** = $p \leq 0.01$, *** = $p \leq 0.001$.

**Supplementary Table 1.** ASO and gRNA Sequences. Items with an * are fully 2'-MOE, all C are $m^5C$, all U are $m^5U$; Halo = chloroalkane HaloTag-ligand connected through 5'-Amino-5 modifier or amino modifier C2-dT (Glen Research). sgRNAs for REPAIRv2 and DECOR are canonical RNA nucleotides. Bolded sequences indicate the direct repeat sequence unique to PspCas13b (REPAIRv2) or RfxCas13d (DECOR).

| Name | Sequence* |
|---|---|
| **Halo-Spinraza*** | 5'-Halo-UCACUUUCAUAAUGCUGG-3' |
| **ACTB ASO 1*** | 5'-Halo-GCAAUGCUAUCACCUCCC-3' |
| **ACTB ASO 2*** | 5'-Halo-AUGCCAAUCUCAUCUUGU-3' |
| **REPAIR ACTB ASO 2 gRNA** | 5'-AUGAGAUUGGCAUGGCUUUAUUUGUUUUUU**GUUGUGGAAGGUCCAGUUUUGAGGGGCUAUUACAAC**-3' |
| **DECOR ACTB ASO 2 gRNA** | 5'-**AACCCCUACCAACUGGUCGGGGUUUGAAAC**AAACCUAACUUGCGCAGAAAACAAGAUGAG-3' |
| **ACTB ASO 3*** | 5'-Halo-CGUAGAUGGGCACAGUGU-3' |
| **ACTB ASO 4*** | 5'-Halo-GCUUCUCCUUAAUGUCAC(dG)-3' |
| **ACTB ASO 5*** | 5'-Halo-CCUUCUGACCCAUGCCCA-3' |
| **ACTB ASO 6*** | 5'-Halo-AUAGCACAGCCUGGAUAG-3' |
| **Halo-Spinraza 2*** | 5'-UCACUUUCA(dT-Halo)AAUGCUGG-3' |
| **4 nt Bulge-forming ASO*** | 5'-UCGAGGCAU(dT-Halo)GCGAUCACU-3' |
| **6 nt Bulge-forming ASO*** | 5'-CUCGAGGCA(dT-Halo)CGAUCACUU-3' |
| **9 nt Bulge-forming ASO*** | 5'-UCUCGAGGC(dT-Halo)GAUCACUUU-3' |
| **PTEN ASO 1*** | 5'-Halo-ACUCAAAGUACAUGAACU-3' |
| **PTEN ASO 2*** | 5'-ACACAGGUAAC(dT-Halo)GGAACUCAAAG-3' |
| **PTEN ASO 3*** | 5'-ACACACAGGUAAC(dT-Halo)GGAACUCAAAGUA-3' |
| **PTEN ASO 4*** | 5'-CACAGGUAACG(dT-Halo)GGGAACUCAAA-3' |
| **REPAIR PTEN gRNA** | 5'**-** AUGUACUUUGAGUUCCCUUAGCCGUUACCU**GUUGUGGAAGGUCCAGUUUUGAGGGGCUAUUACAAC**-3' |
| **DECOR PTEN gRNA** | 5'-**AACCCCUACCAACUGGUCGGGGUUUGAAAC**ACCCACACGACGGGAAGACAAGUUCAUGUA-3' |

**Supplementary Table 2.** ASO thermodynamics. Data was generated from the Oligoscreen web server (https://rna.urmc.rochester.edu/RNAstructureWeb/Servers/oligoscreen/oligoscreen.html) using the RNA setting and 310.15K temperature. The overall duplex energy was determined from the IntaRNA web server (https://rna.informatik.uni-freiburg.de/IntaRNA/Input.jsp).

| Name | dGbimolecular | dGunimolecular | dGduplex | Overall Duplex Energy* | dG2B Pat5' | dG2B Pat3' |
|---|---|---|---|---|---|---|
| **Halo-Spinraza** | -1.8 | 0 | -27.9 | -21.43 | 4.5 | 5.9 |
| **ACTB ASO 1** | -7.4 | 0 | -34.5 | -17.73 | 6 | 6.6 |
| **ACTB ASO 2** | -0.9 | 0 | -28.7 | -20.93 | 2.7 | 4.3 |
| **ACTB ASO 3** | -8.4 | 0 | -34.0 | -25.37 | 5.1 | 4.3 |
| **ACTB ASO 4** | -2.6 | 0 | -36.9 | -20.42 | 5.9 | 4.9 |
| **ACTB ASO 5** | -7.0 | 0 | -32.9 | -29.94 | 2.4 | 3.9 |
| **ACTB ASO 6** | -1.8 | 0 | -32.2 | -21.84 | 6 | 5.1 |
| **Halo-Spinraza2** | -1.8 | 0 | -27.9 | -21.43 | 4.5 | 5.9 |
| **Reporter Bulge ASO 1** | -10.1 | -0.7 | -35.2 | -22.19 | 4.3 | 4.3 |
| **Reporter Bulge ASO 2** | -11.9 | -3.1 | -34.7 | -16.95 | 4.5 | 2.5 |
| **Reporter Bulge ASO 2** | -10.4 | -0.5 | -33.7 | -14.59 | 4.5 | 1.8 |
| **PTEN ASO 1** | -7.0 | 0 | -26.2 | -20.12 | 4.3 | 4.3 |
| **PTEN ASO 2** | -7.2 | -2.7 | -39.5 | -13.36 | 4.3 | 3.5 |
| **PTEN ASO 3** | -8.0 | -2.7 | -46.8 | -16.05 | 4.3 | 3.0 |
| **PTEN ASO 4** | -8.6 | -1.1 | -41.0 | -17.57 | 4.3 | 1.8 |

**Supplementary Table 3.** Primer Sequences

| Name | Sequence |
|---|---|
| **BamHI-8.20-F E. Coli** | 5'-AAAGGATCCGAATTCATGTCTGAGGTGGAGTTTTCCC-3' |
| **BamHI-8.20-R E. Coli** | 5'-AAAGGATCCTCCTCCGCCGTCGGTGGAGCTCTGGG-3' |
| **Downstream GFP F** | 5'-AGGTGAACTTCAAGATCCGC-3' |
| **mCherry-R** | 5'-TTGGTCACCTTCAGCTTGG-3' |
| **ACTB ASO1 F** | 5'-CCTTCTACAATGAGCTGCGTG-3' |
| **ACTB ASO1 R** | 5'-CCATCTCTTGCTCGAAGTCC-3' |
| **ACTB ASO2 F** | 5'-TGGACTTCGAGCAAGAGATGG-3' |
| **ACTB ASO2 R** | 5'-CGATCCACACGGAGTACTTGC-3' |
| **ACTB ASO3 F** | 5'-CCGCAAATGCTTCTAGGCG-3' |
| **ACTB ASO3 R** | 5'-CTCCCCTGTGTGGACTTGG-3' |
| **ACTB ASO4 F** | 5'-GGAGGTGATAGCATTGCTTTCG-3' |
| **ACTB ASO4 R** | 5'-AGGTGTGCACTTTTATTCAACTGG-3' |
| **T7 Actin F** | 5'-TAATACGACTCACTATAGGATGATGATATCGCCGCG-3' |
| **Actin R** | 5'-CTAGAAGCATTTGCGGTGGACG-3' |
| **ACTB 3'UTR 2** | 5'-CAAGTCAGTGTACAGGTAAGCCC-3' |
| **ACTB ASO 5 Seq Pri** | 5'-CTGTGCTATCCCTGTACGCCTCTGG-3' |
| **Spinraza DR GAACA 1** | 5'-GATCCCCAGCATTATGAAAGTGATCGCGAACAATGCCTCGAGAGC-3' |
| **Spinraza DR GAACA 2** | 5'-GGCCGCTCTCGAGGCATTGTTCGCGATCACTTTCATAATGCTGGG-3' |
| **Spinraza DR ATACA 1** | 5'-GATCCCCAGCATTATGAAAGTGATCGCATACAATGCCTCGAGAGC-3' |
| **Spinraza DR ATACA 2** | 5'-GGCCGCTCTCGAGGCATTGTATGCGATCACTTTCATAATGCTGGG-3' |
| **Spinraza DR CTACA 1** | 5'-GATCCCCAGCATTATGAAAGTGATCGCCTACAATGCCTCGAGAGC-3' |
| **Spinraza DR CTACA 2** | 5'-GGCCGCTCTCGAGGCATTGTAGGCGATCACTTTCATAATGCTGGG-3' |
| **Spinraza DR GGAGG 1** | 5'-GATCCCCAGCATTATGAAAGTGATCGCGGAGGATGCCTCGAGAGC-3' |
| **Spinraza DR GGAGG 2** | 5'-GGCCGCTCTCGAGGCATCCTCCGCGATCACTTTCATAATGCTGGG-3' |
| **Spinraza DR GTATG 1** | 5'-GATCCCCAGCATTATGAAAGTGATCGCGTATGATGCCTCGAGAGC-3' |
| **Spinraza DR GTATG 2** | 5'-GGCCGCTCTCGAGGCATCATACGCGATCACTTTCATAATGCTGGG-3' |
| **ACTB CTATG ASO F** | 5'-CACAGAGCCTCGCCTTTGC-3' |
| **ACTB CTATG ASO R** | 5-GCTGGGGTGTTGAAGGTCTC-3' |
| **ACTB GTACG ASO F** | 5'-TGGGCATGGGTCAGAAGG-3' |
| **ACTB GTATG ASO R** | 5'-CCTCAGGGCAGCGGAACC-3' |
| **ACTB TTATT ASO F** | 5'-GGAGATCACTGCCCTGGC-3' |
| **PTEN Q245X F** | 5'-TCTATGGGGAAGTAAGGACC-3' |
| **PTEN Q245X R** | 5'-GTTGGCTTTGTCTTTATTTGC-3' |
| **REPAIR ACTB ASO 2 gRNA 20 nt F** | CACCGAAAAAACAAACAAAGCCATGCCAATCTCAT |
| **REPAIR ACTB ASO 2 gRNA 20 nt R** | CAACATGAGATTGGCATGGCTTTGTTTGTTTTTTC |
| **REPAIR PTEN Q245X gRNA 20 nt F** | CACCGAGGTAACGGCCAAGGGAACTCAAAGTACAT |
| **REPAIR PTEN Q245X gRNA 20 nt R** | CAACATGTACTTTGAGTTCCCTTGGCCGTTACCTC |

**Supplementary Table 4.** Halo-ASO ESI LC-MS Characterization.

| Name | Expected Mass (Da) | Observed Mass (Da) |
|---|---|---|
| **Halo-Spinraza 1** | 7,322.84 | 7,322.86 |
| **ACTB ASO 1** | 7,390.87 | 7,390.92 |
| **ACTB ASO 2** | 7,296.85 | 7,296.89 |
| **ACTB ASO 3** | 7,423.85 | 7,423.89 |
| **ACTB ASO 4** | 7,615.91 | 7,615.93 |
| **ACTB ASO 5** | 7,283.90 | 7,283.95 |
| **ACTB ASO 6** | 7,390.87 | 7,390.92 |
| **Halo-Spinraza 2** | 7,179.82 | 7.179.86 |
| **Bulge ASO 1** | 7,606.93 | 7,606.98 |
| **Bulge ASO 2** | 7,580.94 | 7,581.00 |
| **Bulge ASO 3** | 7,572.91 | 7,572.94 |
| **PTEN ASO 1** | 7,333.88 | 7,333.37 |
| **PTEN ASO 2** | 9,173.33 | 9,173.22 |
| **PTEN ASO 3** | 10,702.70 | 10,702.93 |
| **PTEN ASO 4** | 9,189.32 | 9,189.51 |

**Supplementary Scheme 1.** Synthesis of NHS-Halo ligand (6)[1, 2].

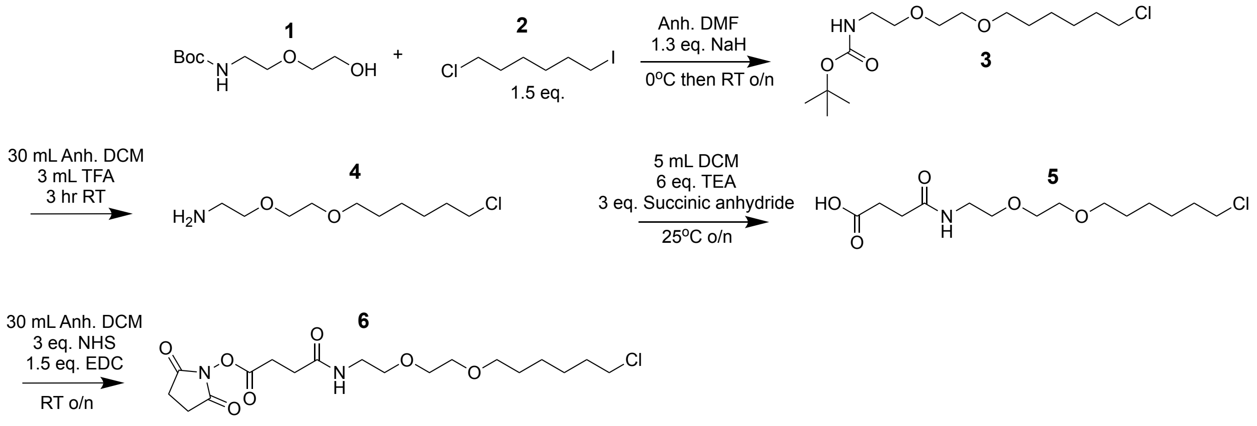
